# Immunopeptidomics informs discovery and delivery of *Mycobacterium tuberculosis* MHC-II antigens for vaccine design

**DOI:** 10.1101/2024.10.02.616386

**Authors:** Owen Leddy, Paul Ogongo, Julia Huffaker, Mingyu Gan, Ryan Milligan, Sheikh Mahmud, Yuko Yuki, Kidist Bobosha, Liya Wassie, Mary Carrington, Qingyun Liu, Joel D. Ernst, Forest M. White, Bryan D. Bryson

## Abstract

No currently licensed vaccine reliably prevents pulmonary tuberculosis (TB), a leading cause of infectious disease mortality. Developing effective new vaccines will require identifying which of the roughly 4000 proteins in the *Mycobacterium tuberculosis* (*Mtb*) proteome are presented on MHC class II (MHC-II) by infected human phagocytes and can be recognized by CD4+ T cells to mediate protective immunity. Vaccines must also elicit T cell responses recognizing the same peptide-MHC complexes presented by infected cells, and successful presentation of target human MHC-II peptides is currently challenging to evaluate and optimize. Here, we define antigenic targets for TB vaccine development by using mass spectrometry (MS) for proteome-wide discovery of *Mtb* epitopes presented on MHC-II by infected human cells. We next iteratively design and evaluate candidate mRNA vaccine immunogens, revealing design principles that enhance presentation of target MHC-II peptides. Our results will inform the development of new TB vaccine candidates.

## Introduction

*Mycobacterium tuberculosis* (*Mtb*) is the causative agent of tuberculosis (TB) and currently causes more deaths per year than any other human pathogen besides SARS-CoV-2.^1^ As of 2022, the annual death toll of TB globally was 1.3 million,^1^ and new interventions to prevent and treat TB are urgently needed. The live attenuated vaccine Bacillus Calmette Guerin (BCG) protects against disseminated forms of TB in children, but provides limited and highly variable protection against pulmonary TB disease in adults.^2,3^ New vaccines with greater efficacy against pulmonary TB in adolescents and adults could prevent millions of deaths and hasten the slow decline in global TB cases.^4^

Antigen-specific CD4+ T cell responses are an essential component of protective immunity to *Mtb* infection.^5–8^ In animal models, CD4+ T cells can only control bacterial replication in infected cells that they directly recognize via MHC-II,^9^ implying that TB vaccines should prime CD4+ T cell responses against target antigens that are presented on MHC-II by cells infected with live *Mtb.* CD4+ T cell reactivity to a range of *Mtb* antigens has been measured^10–12^, but which of the roughly 4000 proteins in the *Mtb* proteome are presented on MHC-II by human phagocytes infected with virulent *Mtb* has not been directly defined. Translation of new TB vaccine candidates to human use also remains a significant challenge. Vaccine candidates tested in model systems that do not include human MHC or do not directly assess presentation of peptides matching those presented in infected phagocytes may prime T cell responses that fail to recognize *Mtb*-infected cells in humans.^13^ Here, we established a pipeline for vaccine target discovery and immunogen design in TB that centers on human antigen presentation biology, combining immunopeptidomics, bacterial genomics, analysis of T cell responses in clinical samples, and iterative design and testing of mRNA-encoded immunogens. These studies identified secreted and cell envelope-associated *Mtb* antigens presented on MHC-II in infected human phagocytes that are immunogenic and highly conserved across global *Mtb* genomic diversity. We also present principles for the design of mRNA vaccine immunogens that are inspired by the cell biology of antigen processing during infection and quantitatively enhance presentation of target epitopes on MHC-II by multiple orders of magnitude.

## Results

### Identification of Mtb antigens presented on MHC-II in infected human phagocytes

To define potential antigenic targets for TB vaccines, we established an MS-based immunopeptidomics workflow to identify *Mtb*-derived peptides presented on MHC-II by phagocytes infected with *Mtb*. We used primary human cells and selected human monocyte-derived dendritic cells (hMDCs) as our *Mtb* infection model, as peptide binding preferences of MHC-II differ between humans and other species and hMDCs presented higher levels of total surface MHC-II than did monocyte-derived macrophages (hMDMs) (Figure 1a-b). Both hMDCs and hMDMs can be infected by *Mtb in vivo*.^14–16^ *Mtb* infection of hMDCs did not significantly alter surface levels of HLA-DR and modestly increased surface levels of HLA-DQ, suggesting that MHC-II antigen presentation was not impaired in infected hMDCs (Figure 1c-d). To identify *Mtb*-derived MHC-II peptides presented by infected cells in a proteome-wide manner, we re-optimized several parameters of our previously established MHC class I immunopeptidomics protocol^17,18^ for MHC-II: the antibody clone and amount used for immunoprecipitation (IP) of MHC-II, the IP duration, the method used to separate MHC-II-associated peptides from MHC-II protein, and offline fractionation of peptides prior to MS analysis (Supplementary figure 1). We also made improvements based on initial results in *Mtb*-infected human phagocytes, such as modifying the gradient used for online liquid chromatography during MS analysis to better separate MHC-II peptides (Supplementary table 1).

**Figure 1.**
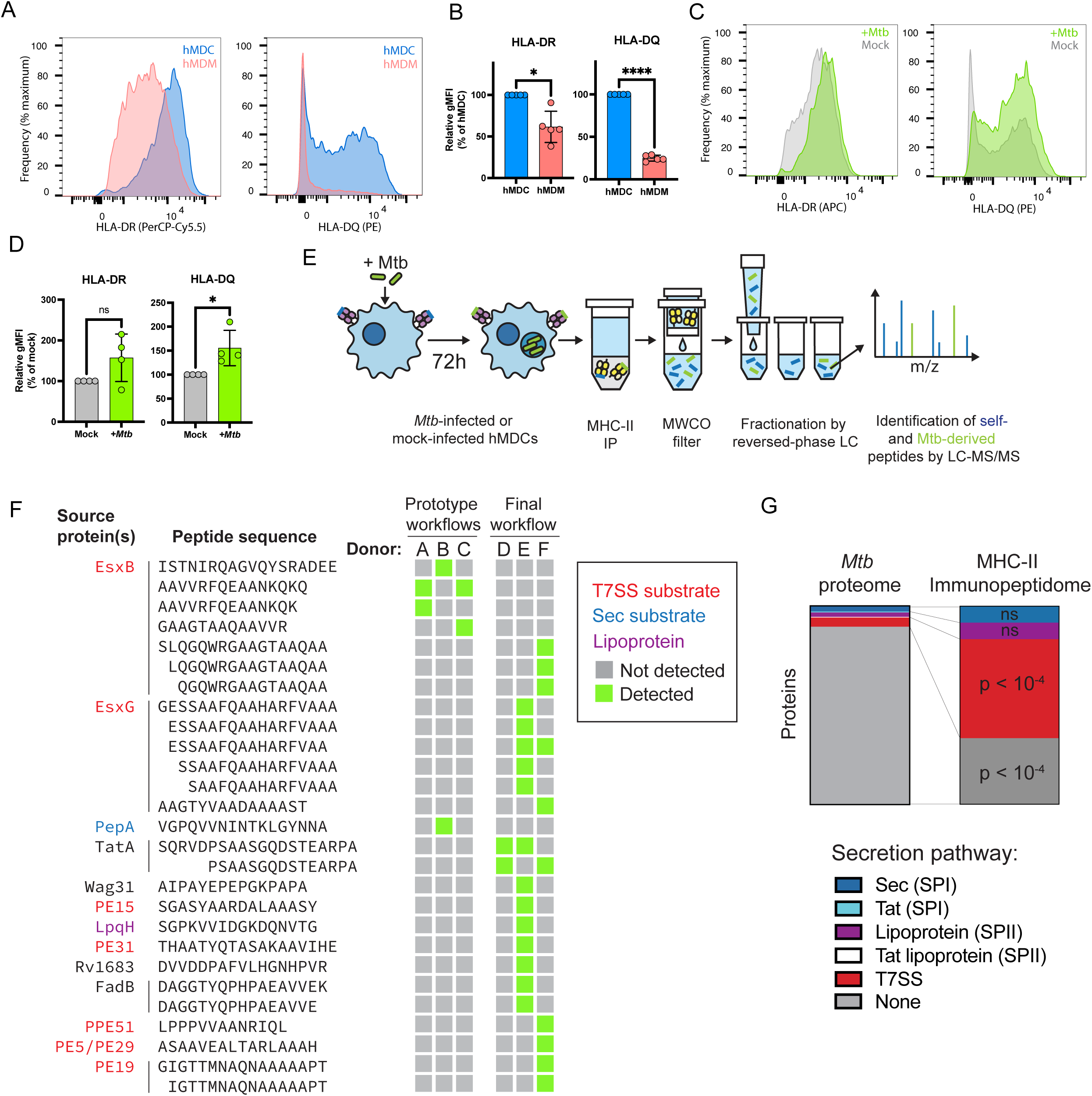
Immunopeptidomics identifies potential vaccine targets presented on MHC-II in *Mtb*-infected human dendritic cells. a) Representative histograms of cell surface HLA-DR and HLA-DQ in hMDMs and hMDCs, as measured by flow cytometry. b) Geometric mean fluorescence intensity of HLA-DR and HLA-DQ cell surface staining in hMDMs and hMDCs determined by flow cytometry (* p < 0.05, **** p < 0.0001 paired ratio t-test; error bars represent standard deviation of n = 5 donors) c) Representative histograms of cell surface HLA-DR and HLA-DQ in *Mtb*-infected and mock-infected hMDCs as measured by flow cytometry 72 hours after infection at MOI 2.5. d) Geometric mean fluorescence intensity of HLA-DR and HLA-DQ cell surface staining in *Mtb*-infected and mock-infected hMDCs as measured by flow cytometry (* p < 0.05, ns = not significant, paired ratio t-test; error bars represent standard deviation of n = 4 donors). e) Schematic outline of final, optimized MHC-II immunopeptidomics workflow (see Methods). f) Peptide sequences and source proteins for *Mtb*-derived MHC-II peptides identified by immunopeptidomics in hMDCs 72 hours after infection at MOI 2.5. Green boxes indicate which donors presented each peptide. g) Enrichment analysis of Mtb peptides presented on MHC-I and their source proteins, categorized by protein secretion pathway using SignalP 6.0^31^ and a curated set of known T7SS substrates. p-values were determined using the hypergeometric test with Bonferroni correction for multiple hypothesis testing (see Methods).

A subset of secreted *Mtb* proteins have been widely used in experimental TB subunit vaccine candidates.^19–23^ These antigens were originally selected in part on the basis of their abundance in *Mtb* culture filtrates and their recognition by T cells in *Mtb*-infected animals,^24^ which may or may not correlate with presentation on MHC-II by infected human cells. We therefore hypothesized that a proteome-wide discovery approach using immunopeptidomics in human cells would reveal additional antigenic targets and help us prioritize among previously described targets. We isolated peptide-MHC-II complexes from *Mtb*-infected or mock-infected hMDCs derived from 6 donors expressing a total of 44 MHC-II variants (unique alpha chain-beta chain allele pairs; Supplementary Table 2) and analyzed the associated peptides by MS. We identified thousands of total MHC-II peptides per analysis (Supplementary table 1), including 27 *Mtb*-derived peptides derived from 13 source proteins (Figure 1f). Peptides derived from 3 source proteins (EsxB, EsxG, and TatA) were presented by hMDCs from multiple donors. Some of these peptides comprise sets of overlapping or nested sequences, as expected given that the ends of the MHC-II binding groove are open and can accommodate peptides with variable flanking sequences.^25^ We validated the identity of at least one peptide per nested set by internal standard parallel reaction monitoring (IS-PRM, also known as SureQuant^26^), comparing the MS/MS spectrum and retention time of each peptide with those of an internal stable isotope labeled (SIL) synthetic standard (Supplementary figure 2). Consistent with previous MHC-II immunopeptidome studies,^27^ a small number of MHC-I peptides (9- to 11-mer peptides matching MHC-I binding motifs) were co-isolated with MHC-II peptides, including *Mtb*-derived peptides derived from EsxB (RADEEQQQAL and AEMKDAATL) that have previously been reported as CD8+ T cell epitopes.^28,29^ These putative MHC-I peptides are excluded in Figure 1. Our results show that immunopeptidomics can identify MHC-II antigens derived from virulent *Mtb* in infected primary human phagocytes. The antigens identified using this approach include at least one commonly targeted by vaccine candidates (EsxB), but also include several other targets not commonly included in previously tested vaccine candidates. Some commonly targeted secreted antigens (such as FbpA, FbpB, and FbpC) were not detected.^19–23^ This dataset provides a new basis for prioritizing antigenic targets for TB vaccines.

We hypothesized that the subcellular localization of *Mtb* antigens within the bacterium could influence accessibility to host MHC-II antigen processing pathways.^30^ We therefore analyzed the predicted protein export pathways of detected MHC-II antigens using SignalP (Figure 1g).^31^ Substrates of *Mtb*’s type VII secretion systems (T7SSs) were overrepresented in the MHC-II repertoire relative to the encoded *Mtb* proteome (p < 10^-4^, binomial test with Bonferroni correction). These included PE/PPE proteins that are found on the surface of the *Mtb* outer membrane or may form heterodimers with known outer membrane proteins, including PPE51^32^, PE19 (binds PPE51^32^), PE15 (binds PPE20^33^), and PE31 (encoded adjacent to PPE60^34^). Lipoproteins and proteins exported via the Sec pathway were neither enriched nor depleted in the MHC-II immunopeptidome. Proteins with no predicted secretion signal were significantly underrepresented in the MHC-II repertoire (p < 10^-4^, binomial test with Bonferroni correction). Most detected MHC-II antigens without predicted secretion signals derived from proteins that are either integral membrane proteins (TatA^35^), are associated with inner membrane protein complexes (Wag31^36,37^), or act on membrane lipids (Rv1683^38^). These results suggest secreted and membrane-associated *Mtb* antigens localized via multiple pathways are preferentially presented on MHC-II relative to cytosolic proteins.

### Mtb-derived MHC-II epitopes are highly conserved and can elicit T cell responses in humans

*Mtb* strains bearing amino acid substitutions in MHC-II epitopes could potentially evade T cell responses induced by vaccines targeting these regions. We therefore analyzed the genomes of 51,229 *Mtb* isolates from a previously collected global dataset and examined whether nonsynonymous mutations in the MHC-II epitopes we identified by MS occurred at ancestral positions, potentially impacting a large number of descendant strains.^39^ We found that amino acid substitutions in these MHC-II epitopes were located in the terminal branches of the phylogeny, each affecting <0.15% of *Mtb* isolates (Fig. 2a; Supplementary data 1). Nonsense mutations upstream of MHC-II epitopes were also rare and affected no more than 6 isolates for most antigens (Supplementary data 2). The sole exception was the mutation Q53stop in PE31, which affects a clade of 429 isolates in lineage 3 (Supplementary data 2). Our results suggest that pre-existing *Mtb* genomic diversity is unlikely to limit the efficacy of vaccines targeting most *Mtb* antigens identified in our analysis of the MHC-II immunopeptidome.

**Figure 2.**
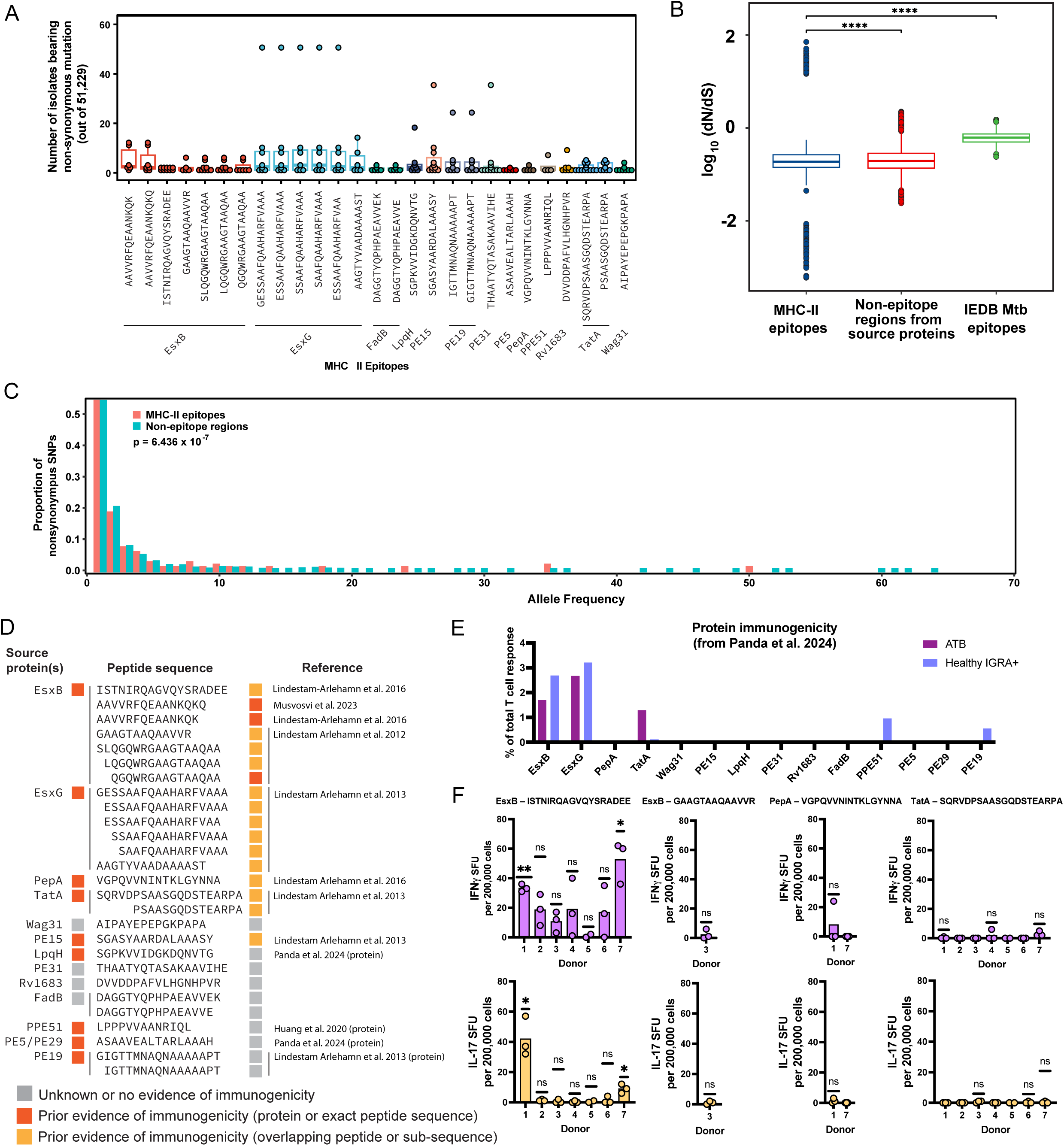
*Mtb* MHC-II antigens identified by immunopeptidomics are highly conserved and can elicit T cell responses *in vivo* in humans. a) Number of isolates bearing nonsynonymous mutations in each MHC-II epitope, out of a database of 51,299 genome sequences of *Mtb* isolates spanning all major lineages.^39^ Each point represents a distinct nonsynonymous mutation occurring in a given epitope. b) Comparison of the dN/dS ratio of sequences encoding *Mtb* MHC-II epitopes with non-epitopes regions of the corresponding gene, as well as with other *Mtb* T cell epitopes reported in IEDB.^40^ (**** p < 0.001, Wilcoxon test.) c) Comparison of the frequency spectrum of nonsynonymous mutations in MHC-II epitopes identified by MS and non-epitope regions of the corresponding genes. p-value determined by Wilcoxon test. d) Schematic outlining evidence for immunogenicity of *Mtb* MHC-II identified by MS.^10–12,42–44^ ^e)^ Contribution of source proteins of *Mtb* MHC-II epitopes identified by MS to human T cell responses in individuals with active TB disease (ATB) or health interferon gamma release assay positive (healthy IGRA+) individuals, as measured by Panda et al. in a prior study.^12^ f) Frequency of IFNɣ and IL-17 spot-forming units (SFUs) in peptide-stimulated PBMCs of healthy IGRA+ individuals, quantified by ELISpot assay. Responses to each peptide were measured in PBMCs from donors expressing the corresponding class II HLA allele (see Supplementary tables 2 and 3). (* p<0.05, ** p<0.01, one-sample t-test.)

We also sought to determine whether MHC-II epitopes show evidence of selective pressure that would favor or disfavor nonsynonymous mutations. MHC-II epitopes identified by MS had slightly but significantly lower pairwise ratios of nonsynonymous to synonymous mutations (dN/dS values) compared to non-epitope regions of the corresponding genes (Figure 2b). Both groups exhibited significantly lower pairwise dN/dS values than other T cell epitopes reported in the immune epitope database (IEDB) from other *Mtb* proteins.^40^ *Mtb* T cell epitopes have previously been reported to be under purifying selection,^41^ which our results suggest is stronger on MHC-II epitopes and antigens we identified by MS than on other reported epitopes. Additionally, the allele frequency spectrum of non-synonymous mutations in MHC-II epitopes was skewed toward lower frequencies relative to the non-epitope regions of the same gene (Figure 2c; p = 4.658 × 10^-16^, pairwise Wilcoxon test), suggesting that the strains carrying nonsynonymous mutations in MHC-II epitopes have fewer descendants than those carrying nonsynonymous mutations in non-epitope regions. Overall, our findings indicate MHC-II epitopes we identified occur in sequences under purifying selective pressure.

Having identified conserved *Mtb*-derived antigenic targets presented on MHC-II *in vitro*, we next asked whether these antigens can be recognized by human T cells *in vivo*, as target antigens must be immunogenic to mediate protection. We first examined prior evidence of *in vivo* human T cell responses to MHC-II epitopes we identified by MS. T cell responses to 3 of the 27 MHC-II peptides we identified have previously been reported in humans with prior *Mtb* exposure (Figure 2d). Responses to a sub-sequence or partially overlapping peptide have been reported for an additional 14 peptides (Figure 2d).^10,42,43^ At the whole-protein level, T cell responses against 9 of the 13 source proteins of *Mtb* MHC-II peptides were previously reported.^10–12,42,44^ Panda et al.^45^ reported that EsxG and EsxB respectively accounted for 3.85% and 2.69% of detected human T cell responses to an *Mtb*-specific peptide pool in asymptomatic interferon gamma release assay positive (IGRA+) individuals, on a similar order of magnitude to the highest single-antigen response reported in the cohort (EspC, 5.82%)^45^ (Figure 2e). Three other source proteins of MHC-II peptides detected in our MS analyses also contributed to T cell responses in either individuals with active TB disease (ATB) or healthy IGRA+ individuals in the same study (Figure 2e).

To determine whether the exact *Mtb* MHC-II peptide sequences identified by MS could elicit T cell responses *in vivo* in humans, we used an ELISpot assay to measure production of IFNγ and IL-17 elicited by four such peptides in peripheral blood mononuclear cells (PBMCs) of a small cohort of individuals with prior *Mtb* exposure (i.e., IGRA+ individuals) who expressed HLA alleles associated with each epitope (Supplementary table 3). An EsxB-derived MHC-II peptide (ISTNIRQAGVQYSRADEE) stimulated IFNγ-producing T cell responses with an average frequency of >10 spot-forming units (SFU) per 200,000 PBMCs in 6 out of 7 HLA-DQB1*03:01-expressing donors tested, and for 2 donors the IFNγ and IL-17 responses were both statistically significant (p < 0.05, one-sample t test) (Figure 2f). These results show that MHC-II peptides identified by MS can be immunogenic *in vivo* in humans with prior *Mtb* exposure. The three other peptides tested (derived from EsxB, PepA, and TatA respectively) did not stimulate significant production of IFNγ or IL-17 in the donors assayed, suggesting that immunopeptidomics may also capture MHC-II peptides that are not immunodominant during infection. Sequence identity with human proteins is a determinant of immunogenicity that may be important to consider in selecting antigenic targets for TB vaccines. A BLAST search of TatA_64-83_ (SQRVDPSAASGQDSTEARPA) and EsxB_72-89_ (ISTNIRQAGVQYSRADEE) against the human proteome revealed that TatA_64-83_ contains regions of 9 to 11 amino acids with >80% identity to multiple human proteins, whereas EsxB_72-89_ had no significant identity to human proteins (Supplementary figure 3). The lower immunogenicity of TatA_64-83_ therefore could potentially be explained by tolerance due to cross-reactivity with self epitopes.^46,47^

### Quantitative targeted MS enables evaluation and optimization of vaccine immunogens

Whereas measuring the ability of vaccines to drive presentation of human MHC-II epitopes matching those presented by *Mtb*-infected cells would typically require an immunogenicity study in HLA-transgenic mice or a human clinical trial, we established an approach to quickly evaluate vaccine candidates using quantitative targeted MHC-II immunopeptidomics. We reasoned that since MHC-II peptides identified by MS *in vitro* show evidence of immunogenicity following *Mtb* exposure *in vivo*, MS could also provide a rapid *in vitro* assay to test and optimize MHC-II antigen presentation upon vaccine delivery. Our method, based on SureQuant,^26^ is analogous to one successfully used in the context of MHC class I to optimize T cell priming vaccines against SARS-CoV-2.^48^ This approach allowed us to both evaluate the ability of the existing BCG vaccine to drive presentation of MHC-II epitopes matching those presented during *Mtb* infection and optimize the design of novel mRNA-encoded vaccine immunogens, leveraging the programmability of the mRNA platform for rapid design-build-test cycles.

It is not currently known whether antigen presenting cells infected with the BCG vaccine strain present the same peptides on MHC-II as those infected with *Mtb*, and opinions in the field differ as to whether new TB vaccines should aim to boost pre-existing T cell responses elicited by BCG or prime *de novo* responses against an orthogonal set of antigens.^49,50^ Despite the limited efficacy of intradermal BCG vaccination against pulmonary TB in humans, studies in non-human primates have shown that BCG immunization can confer sterilizing protection in a CD4+ T cell-dependent manner when administered intravenously, suggesting MHC-II epitopes presented by both BCG- and *Mtb*-infected cells may be protective.^8,51^ Using SureQuant, we found that BCG-infected HLA-DRB1*01:01-expressing hMDCs presented MHC-II peptides derived from EsxG, PE5/PE29, and PE19 that matched those presented by *Mtb*-infected cells (Figure 3a-d). As expected, BCG-infected hMDCs did not present peptides derived from EsxB, which is encoded in the region of difference 1 (RD1) locus that is present in *Mtb* but deleted in BCG^52^ (Figure 3e,f). All non-RD1 *Mtb* MHC-II epitopes we identified and their source proteins are highly conserved in BCG (Supplementary table 4). These results show that BCG can drive presentation of non-RD1 epitopes conserved in *Mtb*, which may inform the design of new vaccines to boost CD4+ T cell responses elicited by BCG.

**Figure 3.**
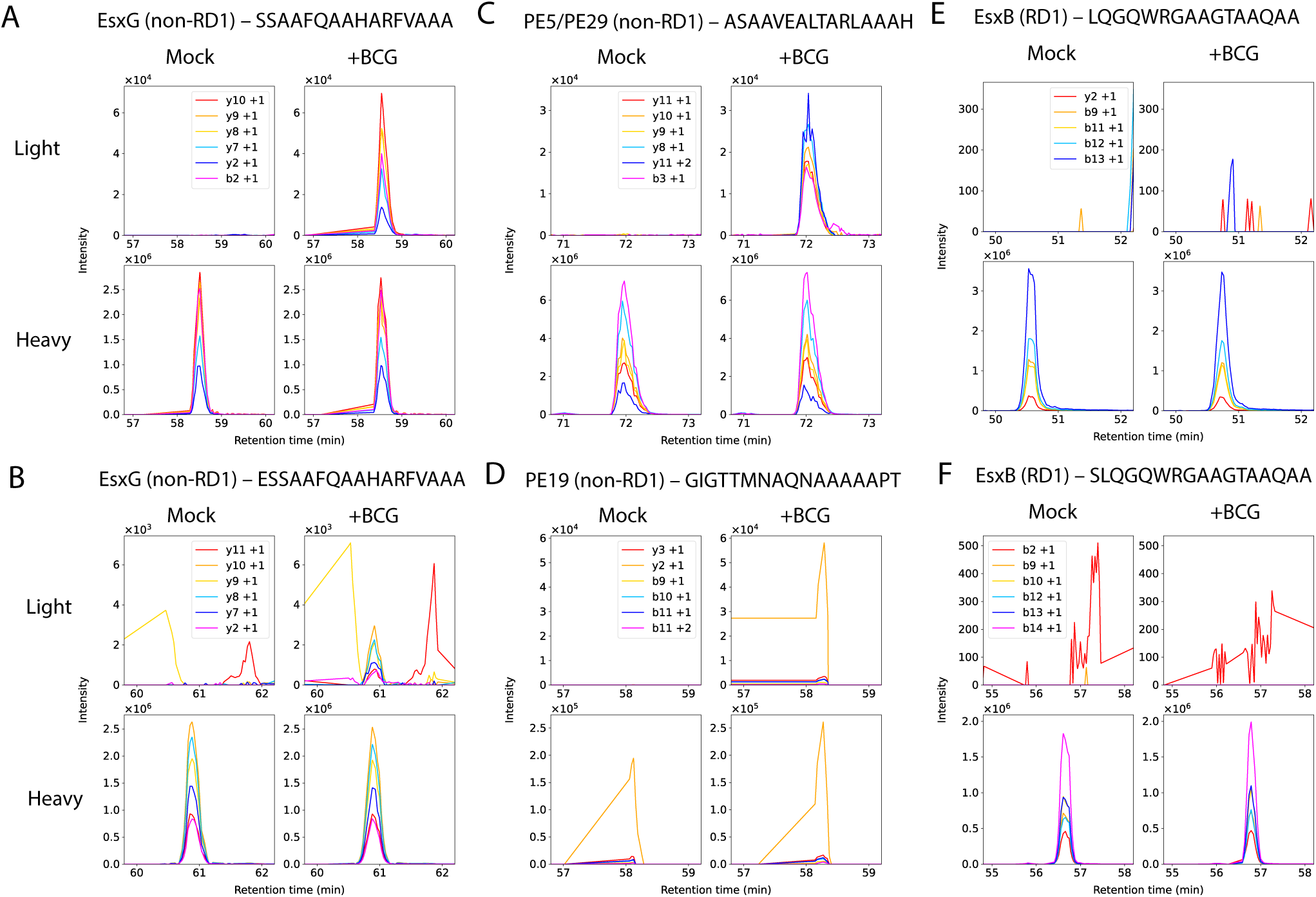
BCG-infected hMDCs present conserved non-RD1 antigens on MHC-II. SureQuant fragment ion chromatograms demonstrating detection of MHC-II peptides derived from EsxG (a,b), PE5/PE29 (c), and PE19 (d) in BCG-infected, HLA-DRB1*01:01-expressing hMDCs (24 hours after infection at MOI 2.5) but not EsxB-derived peptides (e,f). Representative of n = 3 donors.

We next applied our targeted MS approach to optimize candidate mRNA vaccines, which are a promising tool for priming T cell responses^53^ but have not yet been extensively optimized for use against TB. We hypothesized that optimal presentation of *Mtb* MHC-II peptides would require drawing inspiration from the cell biology of *Mtb* antigen processing during infection in order to deliver mRNA-encoded immunogens in a manner that recapitulates the same antigen processing steps. In infected hMDCs, a subset of *Mtb*-containing phagosomes co-localize with the late endosome and lysosome markers LAMP1 and Cathepsin B (CTSB) (Supplementary figure 4a-b), leading us to investigate the contribution of endolysosomal pathways of antigen processing using SureQuant. Treating infected hMDCs with E64d to inhibit cysteine cathepsins did not affect presentation of *Mtb* MHC-II antigens, suggesting that other proteases may have a predominant or redundant role in processing *Mtb* MHC-II peptides (Supplementary figure 4c). We hypothesized that autophagy might contribute to processing of *Mtb* MHC-II antigens, as autophagy has been reported to direct *Mtb* to autophagolysosomes^54^ and promotes presentation of MHC-II peptides derived from other phagocytic cargos.^55,56^ Treating infected hMDCs with wortmannin to block autophagy via inhibition of phosphoinositide-3 kinase (PI3K) resulted in a lower average level of *Mtb* MHC-II peptide presentation, but this trend was not statistically significant and moreover could be due to any of several pleiotropic effects of PI3K inhibition^57^ (Supplementary figure 4c). In contrast to these other perturbations, inhibiting endomembrane compartment acidification by treating *Mtb*-infected hMDCs with bafilomycin caused a statistically significant decrease in *Mtb* MHC-II antigen presentation (Supplementary figure 4c; p < 0.05, two-way main effects ANOVA), suggesting that localization of *Mtb* antigens to acidified endomembrane compartments such as lysosomes may contribute to processing of *Mtb* MHC-II peptides. These results informed our approach to optimizing mRNA-encoded immunogens.

To determine whether delivery of mRNA-encoded *Mtb* antigens to lysosomes could enhance presentation of target MHC-II peptides by mimicking antigen processing during infection, we produced nucleoside-modified mRNAs encoding EsxB, EsxG, or EsxA translationally fused to signals targeting them to a range of subcellular compartments: namely the cytosol, mitochondria, endoplasmic reticulum (ER), endosomes, or lysosomes (Figure 4a, Supplementary table 5; see Methods). HLA-DRB1*01:01-expressing hMDCs transfected with EsxG- and EsxB-encoding mRNAs presented higher levels of target EsxG-derived MHC-II epitopes when antigens were trafficked to the lysosome, relative to other subcellular compartments (Figure 4b,c). These results show that delivering mRNA-encoded *Mtb* antigens to lysosomes can increase presentation of MHC-II peptides that match those presented in infected cells.

**Figure 4.**
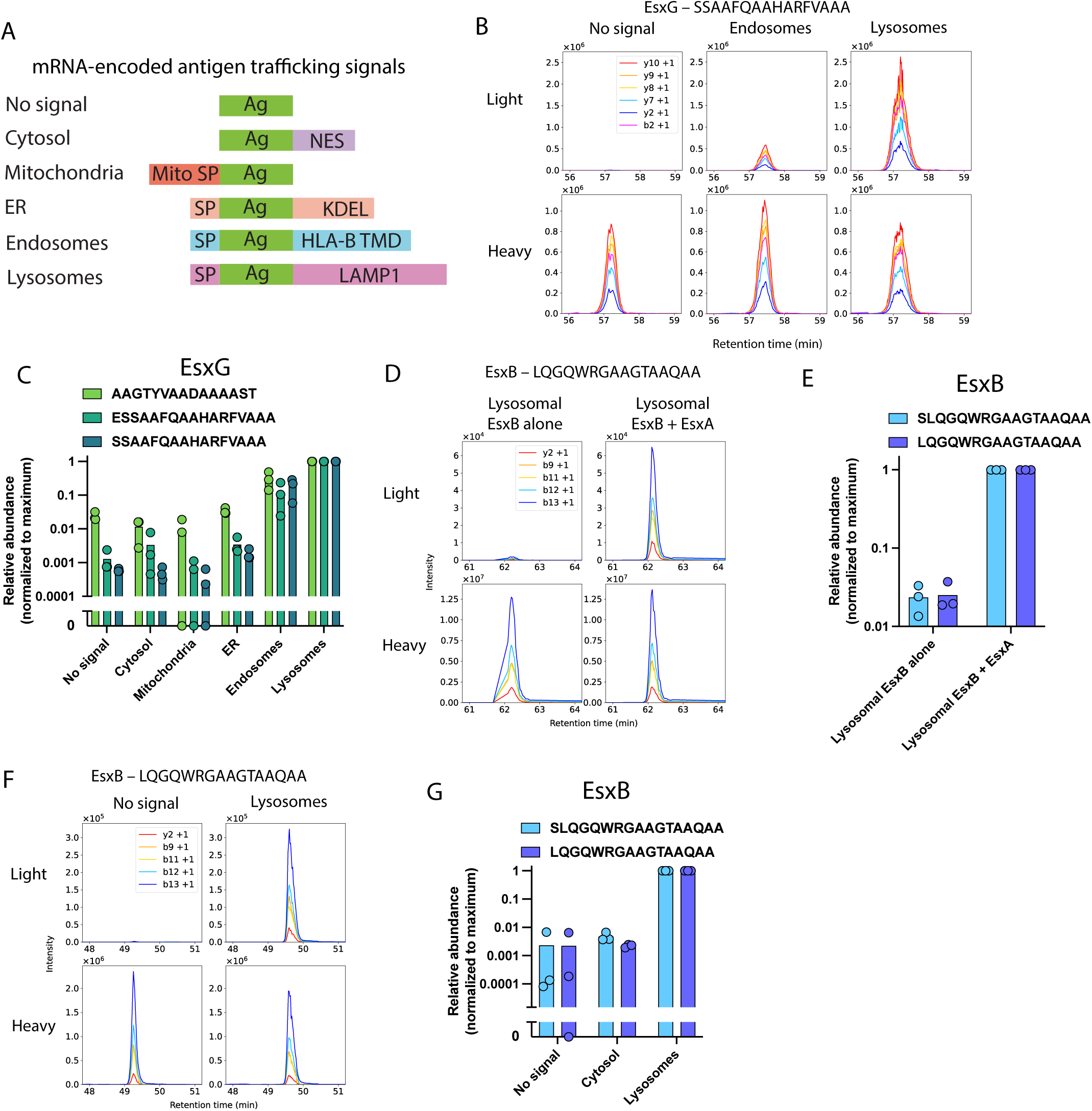
Quantitative targeted MS reveals design principles for mRNA-encoded immunogens that optimize presentation of target *Mtb* antigens on MHC-II. a) Schematic representing fusion protein designs for targeting mRNA-encoded *Mtb* antigens to different subcellular compartments. (Ag = antigen; HLA-B TMD = human leukocyte antigen B transmembrane domain; SP = signal peptide, i.e., N-terminal secretion signal sequence; Mito SP = mitochondrial import signal peptide; NES = nuclear export signal, used to mitigate possible coincidental nuclear import signals in heterologous proteins expressed in mammalian cells.^91^) b) Representative SureQuant fragment ion chromatograms for an EsxG-derived MHC-II epitope in HLA-DRB1*01:01-expressing hMDCs transfected with mRNAs encoding EsxG and EsxB with no subcellular localization signal, or endosomal or lysosomal targeting signals. c) Quantification of EsxG-derived MHC-II peptides presented by HLA-DRB1*01:01-expressing hMDCs transfected with mRNAs encoding EsxG and EsxB with no subcellular localization signal, or cytosolic, mitochondrial, ER, endosomal, or lysosomal targeting signals for n = 3 donors. d) Representative SureQuant fragment ion chromatograms for EsxB-derived MHC-II peptides presented by HLA-DRB1*01:01-expressing hMDCs transfected with mRNAs encoding lysosome-targeted EsxB alone or co-transfected with lysosome-targeted EsxA and EsxB. e) Quantification of EsxB-derived MHC-II peptides presented by HLA-DRB1*01:01-expressing hMDCs transfected with mRNAs encoding lysosome-targeted EsxB alone or co-transfected with lysosome-targeted EsxA and EsxB, for n = 3 donors. f) Representative SureQuant fragment ion chromatograms for an EsxB-derived MHC-II epitope in HLA-DRB1*01:01-expressing hMDCs transfected with mRNAs encoding EsxB and EsxA with no subcellular localization signal or with lysosomal targeting signals. g) Quantification of EsxB-derived MHC-II peptides presented by HLA-DRB1*01:01-expressing hMDCs transfected with mRNAs encoding EsxA and EsxB with no subcellular localization signal, or cytosolic or lysosomal targeting signals for n = 3 donors.

In contrast to EsxG, trafficking of EsxB to lysosomes was not sufficient to drive efficient presentation of target EsxB MHC-II peptides when EsxB was delivered alone or in combination with EsxG (Supplementary Figure 5), suggesting a need for additional strategies to more closely mimic antigen processing that occurs during infection. In the context of *Mtb* infection, EsxB binds to EsxA, and the pair is secreted by the ESX-1 type VII secretion system as a heterodimer that can remain stably associated under acidic conditions.^58,59^ We hypothesized that EsxA and EsxB might need to be delivered together to recapitulate the sequence of antigen processing steps that produces EsxB-derived MHC-II peptides during infection. Co-transfection of mRNAs encoding lysosome-localized EsxB and EsxA led to efficient presentation of target EsxB MHC-II peptides, whereas delivery of mRNA encoding lysosome-localized EsxB alone did not (Figure 4d,e). hMDCs efficiently presented target EsxB MHC-II peptides when EsxB and EsxA were directed to the lysosome, but not when both were directed to the cytosol (Figure 4f,g), showing that directing mRNA-encoded antigens to the lysosome enhances presentation of target EsxB peptides on MHC-II, just as it does for EsxG peptides. These results suggest that interactions between heterodimeric pairs of *Mtb* proteins may influence processing of some MHC-II antigens, and the need to co-deliver heterodimeric antigens to enhance MHC-II peptide presentation may have to be empirically assessed for individual antigens or epitopes of interest.

## Discussion

In this study, we sought to address major hurdles in human TB vaccine development by establishing an approach to rapidly identify and characterize candidate antigens and quantitatively evaluate vaccine platforms that deliver those antigens. *Mtb* antigens presented on MHC-II primarily derived from secreted and cell-envelope-associated proteins. These epitopes are highly conserved across *Mtb* lineages, and many are immunogenic in humans. Furthermore, some of these epitopes derive from essential proteins that are unlikely to be deleted under vaccine selective pressure (e.g., EsxG^60^, TatA^61^, and PE5^60^). Notably, we identified one MHC-II peptide derived from PepA, one of two antigens included in a leading TB subunit vaccine candidate.^62^ One EsxB-derived peptide we identified is the cognate epitope of a TCR specificity group associated with a lower risk of progression to TB disease,^43^ showing that epitopes identified by MS can generate protective T cell responses.

We analyzed sequence variation in *Mtb* MHC-II epitopes presented by infected cells across global *Mtb* genomic diversity and found that these epitopes were highly conserved in all major *Mtb* lineages and potentially subject to purifying selection that disfavors mutations that could disrupt T cell recognition. *Mtb* T cell epitopes under purifying selection have previously been described as potential “decoy” antigens that might be conserved because T cell responses against these epitopes are adaptive for *Mtb*.^41,63^ An alternate explanation – equally consistent with our data – could be that *Mtb* is under immune selective pressure to eliminate MHC-II epitopes that are presented by infected cells, leaving only epitopes that occur in regions that are essential for fitness and therefore highly conserved. A previously identified protective epitope^43^ that also appeared in our MS analyses (sequence AAVVRFQEAANKQKQ) is also highly conserved, suggesting that epitopes with a dN/dS ratio indicating purifying selection can be protective T cell antigens and may not necessarily be immunological decoys.

Many *Mtb* MHC-II antigens we identified elicit T cell responses in humans with prior *Mtb* exposure, validating the *in vivo* relevance of our findings. We also identified novel MHC-II peptides that did not elicit production of IFNγ or IL-17 in PBMCs of HLA-matched donors we assayed. This may be attributable to sample size and/or tolerance to antigens with partial sequence identity to self proteins. If the differences in observed T cell responses instead reflect an intrinsic difference in immunodominance among epitopes, then a vaccine could potentially prime a T cell response to epitopes that would otherwise be subdominant during *Mtb* infection, increasing the breadth of the T cell response to infection. By identifying non-immunodominant targets, MS data may complement information obtained through T cell assays. In the setting of cancer, vaccination can prime protective T cell responses against antigens that are otherwise subdominant,^64^ and the same may be true in *Mtb* infection.

In addition to target discovery, we establish a MS-based approach to rapidly evaluate and optimize the ability of vaccines to drive presentation of target *Mtb* MHC-II antigens. BCG can confer sterilizing CD4+ T cell-dependent protection against *Mtb* infection when administered intravenously in non-human primates,^8,51^ but it was not clear *a priori* which *Mtb* MHC-II peptides are also presented by cells infected with BCG. Phagosome maturation and other aspects of the host cell response differ between BCG-infected and *Mtb*-infected cells, which could in principle alter antigen processing.^65,66^ We showed that conserved MHC-II epitopes derived from EsxG, PE19, and PE5/PE29 can be presented by both BCG- and *Mtb*-infected cells. MHC-II epitopes conserved in BCG and *Mtb* could be useful targets for vaccine strategies designed to boost pre-existing T cell responses induced by BCG.^50^

mRNA vaccines can elicit strong T cell responses,^53^ making them a promising platform for future TB vaccine development. To date, however, mRNA vaccines have primarily been deployed against viral infections, and mRNA vaccines targeting intracellular bacterial infections such as TB have not yet been systematically optimized. To establish design principles for TB mRNA vaccines, we compared the ability of mRNA-encoded immunogens directed to different subcellular compartments^67,68^ to drive presentation of MHC-II epitopes matching those presented by *Mtb*-infected cells. Translationally fusing mRNA-encoded antigens to signal sequences that direct them to endomembrane compartments – particularly lysosomes – enabled more efficient presentation of target *Mtb* MHC-II peptides. Directing antigens to lysosomes may also be a useful strategy for immunization against other microbial infections in which CD4+ T cells have a protective role.^69,70^ Increasing the amount of MHC-II peptide presented per amount of mRNA input using this strategy may reduce the dose of mRNA vaccine required.

Both CD4+ and CD8+ T cells can contribute to immune control of *Mtb* infection,^71,72^ so effective TB vaccine designs may need to drive presentation of MHC-I epitopes matching those presented by infected cells as well as MHC-II epitopes. Processing and presentation of *Mtb* MHC-I antigens requires access to the host cell cytosol.^18,73^ Endolysosomal localization signals that improve presentation of target *Mtb* epitopes on MHC-II therefore may not be optimal for presentation of MHC-I epitopes. Nucleic acid vaccines intended to prime both CD4+ and CD8+ T cell responses may need to direct antigens to distinct compartments.

Many *Mtb* MHC-II antigens we identified are T7SS substrates that are secreted along with a binding partner as a heterodimer.^58,74–76^ We detected higher levels of MHC-II peptides derived from mRNA-encoded EsxB when codelivered with its binding partner^58^ EsxA. Protein tertiary structure has previously been shown to influence MHC-II antigen processing,^77^ and protein-protein interactions may similarly affect the accessibility of specific regions of an antigen to proteases. Intriguingly, presentation of detectable levels of EsxG on MHC-II did not require coexpression of its binding partner EsxH, so we propose that antigen codelivery may need to be empirically assessed for individual antigens.

Some commonly targeted *Mtb* antigens included in TB vaccine candidates, such as members of the antigen 85 family (FbpA, FbpB, FbpC), were not detected in our analysis of the MHC-II immunopeptidome. Similarly, a peptide library screen of humans with prior Mtb exposure previously revealed that these antigens comprise a relatively small fraction of the overall human T cell response to *Mtb* antigens.^12^ These previously described antigenic targets may in fact not be efficiently presented on MHC-II in human cells infected with live *Mtb*, but it is also possible they may not have been detected in our MS analyses for other biological or technical reasons. MS analyses have a limit of detection, and data-dependent MS cannot specifically select precursor ions derived from *Mtb* peptides. The dynamics of host-*Mtb* interactions are complex due to differences in human and *Mtb* genotype, bacterial state, and *Mtb*-phagocyte interactions. Here, we establish a modular approach to antigen discovery that can generalize to other experimental conditions of interest, including phagocyte activation, *Mtb* strains, *Mtb* state, and human HLA genotype. These experiments may identify additional antigens that can be presented on MHC-II.

Our results represent the first dataset directly examining the composition of the *Mtb*-derived MHC-II peptide repertoire in infected cells, providing new evidence to inform the prioritization of antigenic targets for TB vaccines. Additional evidence from bacterial genomics and T cell response measurements supports the use of MHC-II antigens identified by MS as vaccine targets, as they are conserved and many can elicit *in vivo* human T cell responses. We also demonstrate that this knowledge can be translated into new vaccine immunogen designs. Establishing a quantitative MS assay for presentation of *Mtb* MHC-II peptides matching those presented by infected cells lets us take advantage of the programmability and flexibility of mRNA as a vaccine delivery platform, enabling iterative design and testing of candidate vaccine immunogens. Using this approach, we demonstrate new immunogen design principles inspired by the cell biology of *Mtb* antigen processing in infected cells that can greatly increase presentation of target *Mtb* MHC-II peptides upon mRNA delivery to human antigen presenting cells. These results will inform the design of vaccine candidates to prime CD4+ T cell responses against *Mtb*.

## Methods

### Mycobacterial culture

*Mycobacterium tuberculosis* (*Mtb*) H37Rv or *Mycobacterium bovis* BCG Pasteur (BCG) was grown in Difco Middlebrook 7H9 media supplemented with 10% OADC, 0.2% glycerol, and 0.05% Tween-80 to mid-log phase.

### Primary cell isolation, differentiation, and culture

For immunopeptidomics, flow cytometry, and microscopy experiments, deidentified buffy coats were obtained from Massachusetts General Hospital. Samples are acquired and provided to research groups with no identifying information. For SureQuant experiments requiring HLA-DRB1*01:01+ cells, leukapheresis samples from HLA typed donors were obtained from StemCell. PBMCs were isolated by density-based centrifugation using Ficoll (GE Healthcare). CD14+ monocytes were isolated from PBMCs using a CD14 positive-selection kit (StemCell). Isolated monocytes were differentiated into hMDCs in R10 media [RPMI 1640 without phenol red (Gibco) supplemented with 10% heat-inactivated FBS (Gibco), 1% HEPES (Corning), 1% L-glutamine (Sigma)] supplemented with 25 ng/mL GM-CSF (Biolegend, 572902) and 25 ng/mL IL-4 (Biolegend, 572902). Media was replaced with fresh cytokine-containing media every 3 days. Monocytes were differentiated into hMDMs in R10 media supplemented with 25 ng/mL M-CSF (Biolegend, 574804) or, where indicated, 25 ng/mL GM-CSF for 6 days. hMDCs were differentiated in tissue culture treated flasks or plates appropriate for a given experiment (VWR), whereas hMDMs were differentiated in ultra-low-attachment plates or flasks in a format appropriate for a given experiment (Corning).

### Raji cell culture

Raji cells were cultured in T75 flasks (VWR) in RPMI 1640 with 10% heat-inactivated FBS (Gibco).

### HLA genotyping

Genomic DNA was extracted from 5x10^6^ PBMCs using a Qiagen DNeasy kit. HLA typing was performed using a targeted next generation sequencing (NGS) method. Briefly, locus-specific primers were used to amplify a total of 26 polymorphic exons of HLA-A & B (exons 1–4), C (exons 1–5), E (exon 3), DPA1 (exon 2), DPB1 (exons 2–4), DQA1 (exon 1–3), DQB1 (exons 2 & 3), DRB1 (exons 2 & 3), and DRB3/4/5 (exon 2) genes with Standard BioTools Access Array system (Standard BioTools, South San Francisco, CA 94080 USA). The 26 Standard BioTools PCR amplicons were harvested from Standard BioTools Access Allay IFC and pooled. Quality and quantity were checked using a Caliper LabChip GX Touch HT Nucleic Acid Analyzer (PerkinElmer, Waltham, MA 02452 USA). The PCR product library was quantitated and subjected to sequencing on an Illumina MiSeq sequencer (Illumina, San Diego, CA 92122 USA). HLA alleles and genotypes were called using the Omixon HLA Explore (version 2.0.0) software (Omixon Biocomputing Ltd., Budapest, Hungary).

### M. tuberculosis and *BCG infection*

The *Mtb* culture was pelleted by centrifugation, washed once with PBS, resuspended in R10 media and centrifuged at low speed (500 rpm for 5 minutes) to pellet clumps, leaving a uniform suspension of bacteria in the supernatant. For MHC-II target discovery experiments, 50 million hMDCs were infected at MOI 2.5 for 4 hr and then washed with PBS to remove extracellular *Mtb*. Infected hMDCs were cultured in R10 media for 72 hours before harvesting. For SureQuant experiments with drug treatment, 10 million hMDCs per condition were infected at MOI 2.5 for 4 hr in the presence of drug (except wortmannin, which is known to inhibit phagocytosis^78^ and was added only after washing away extracellular bacteria with PBS). Final drug concentrations in media were 250 nM for wortmannin, 20 µM for E64d, and 10 nM for bafilomycin. After washing away remaining extracellular bacteria, fresh R10 media containing drug was added and cells were harvested 24 hours after infection. BCG infections were performed analogously.

### In vitro transcription

Nucleoside-modified mRNAs were produced from 1 μg of linearized template DNA using the HiScribe T7 mRNA Kit with CleanCap Reagent (New England Biolabs) according to manufacturer instructions with the following modifications: Cytidine triphosphate and uracil triphosphate were replaced with 5-methyl-cytidine triphosphate and 5-methoxyuridine triphosphate (Trilink). mRNAs were purified using the Monarch RNA Cleanup Kit (New England Biolabs).

### mRNA transfection

For each flask of 1 × 10^7^ hMDCs, 15 μL of MessengerMax lipofectamine (Thermo) was mixed with 125 μL Opti-MEM (Gibco), vortexed for 3 seconds, and incubated for 10 minutes at room temperature. 10 pmol of mRNA was diluted in 125 μL of Opti-MEM, added to the lipofectamine/Opti-MEM mixture (for a final volume 250 μL of Opti-MEM) and incubated at room temperature for an additional 5 minutes. The mixture was then added to the media of the flask of hMDCs. 4 hours after adding the mRNA/lipofectamine/Opti-MEM mixture, the media was removed, the cells were washed with PBS, and fresh R10 media was added. hMDCs were incubated overnight (16-18 hours) after mRNA transfection before being harvested for MHC-II peptide isolation (see below).

### MHC immunoprecipitation

hMDCs were harvested by collecting the culture media (containing any cells in suspension), washing adherent cells with PBS, incubating remaining adherent cells with PBS supplemented with 4 mM EDTA for 15 minutes at 37 °C, gently scraping with a cell scraper, collecting the detached cells, and washing the flask with PBS and collecting the wash. Harvested cells were pelleted by centrifugation. For optimization experiments, Raji cells, which grow in suspension, were collected by centrifugation without additional treatment. The harvested cells were then washed with PBS, and lysed in 1 mL of MHC lysis buffer [20 mM Tris, 150 mM sodium chloride, pH 8.0, supplemented with 1% CHAPS, 1 x HALT protease and phosphatase inhibitor cocktail (Pierce), and 0.2 mM phenylmethylsulfonyl fluoride (Sigma-Aldrich)].

BSL3 protocol: Lysate from *Mtb*-infected, BCG-infected, or mock-infected cells was sonicated using a Q500 ultrasonic bath sonicator (Qsonica) in five 30-second pulses at an amplitude of 60%, cleared by centrifugation at 16,000 x g for 5 minutes, and sterile filtered twice using 0.2 μm filter cartridges (Pall NanoSep).

BSL2 protocol: Lysate from Raji cells or mRNA-transfected hMDCs was sonicated using a VCX-130 probe ultrasonic processor (Sonics) in three 10-second pulses at an amplitude of 30% and cleared by centrifugation at 16,000 x g for 5 minutes.

For quantitative SureQuant experiments (see below), lysate protein concentrations were normalized by BCA assay and 100 fmol of each of three soluble peptide-DRA1*01:01-DRB1*07:01 complexes containing SIL peptides (heavy isotope-labeled peptide-MHCs – hipMHCs) (ImmunAware custom order) were spiked into each sample to be used as internal standards.

Lysates were then added to protein A sepharose beads pre-conjugated with pan-MHC-II antibody (clone Tü39) – or other antibody, where specified – prepared as previously described.^17^ For untargeted discovery MS experiments, 0.25 mg of antibody was used. For targeted MS experiments, 0.1 mg of antibody was used. Beads were incubated with lysate rotating at 4 °C overnight (12–14 hours). Beads were then washed and peptide-MHC complexes eluted as previously described.^17,18^

### MHC-II peptide isolation

MHC-II-associated peptides were purified using 10 kDa molecular weight cutoff filters (Pall NanoSep) as previously described,^17^ snap-frozen in liquid nitrogen, and lyophilized. Where indicated in optimization experiments (Supplementary figure 1) peptides were isolated by solid phase extraction using a C18 spintip (Protea) as previously described^18^ or a column packed in-house with 9 mg of restricted access material solid phase extraction (RAM-SPE) medium from a Shim-pack MAYI ODS cartridge (Shimadzu), following the protocol specified by Bernhardt et al.^79^

### Offline fractionation

Lyophilized peptides were resuspended in solvent A2 (10 mM triethyl ammonium bicarbonate [TEAB], pH 8.0) and loaded onto a fractionation column (a 200 μm inner diameter fused silica capillary packed in-house with 10 cm of 10 μM C18 beads). Peptides were fractionated using an Agilent 1100 series liquid chromatograph using buffers A2 (10 mM TEAB, pH 8.0) and B2 (99% acetonitrile, 10 mM TEAB, pH 8.0). The fractionation column was washed with solvent A2, and peptides were separated using a gradient of 1–5% solvent B2 over 5 min, 5–40% over 60 min, 40–70% over 10 min, hold for 9 min, and 70%–1% over 1 min. Ninety-second fractions were collected, concatenated into 12 tubes, over 90 min in total. Fractions were flash-frozen in liquid nitrogen and lyophilized.

### DDA MS analyses

MHC-I peptide samples were resuspended in 0.1% formic acid. 25% of the sample was refrozen and reserved for later SureQuant validation analyses, while 75% of the sample was used for DDA analysis. For all MS analyses, samples were analyzed using an Orbitrap Exploris 480 mass spectrometer (Thermo Fisher Scientific) coupled with an UltiMate 3000 RSLC Nano LC system (Dionex), Nanospray Flex ion source (Thermo Fisher Scientific), and column oven heater (Sonation). The MHC peptide sample was loaded onto a fused silica capillary chromatography column with an integrated electrospray tip (∼1 μm orifice) prepared and packed in-house with 10 cm of 1.9 μm C18 beads (ReproSil-Pur).

Standard mass spectrometry parameters were as follows: spray voltage, 2.0 kV; no sheath or auxiliary gas flow; ion transfer tube temperature, 275 °C. The Orbitrap Exploris 480 mass spectrometer was operated in data dependent acquisition (DDA) mode. Peptides were eluted using one of two gradients using 0.1% formic acid as buffer A and 70% Acetonitrile, 0.1% formic acid as buffer B:

Gradient 1 (original MHC gradient): 6–25% buffer B over 75 min, 25–45% over 5 min, 45–97% over 5 min, hold for 1 min, and 97% to 3% over 2 min.

Gradient 2 (MHC-II optimized gradient): 6–12% buffer B over 20 min, 12–20% over 45 min, 20– 25% over 10 min, 25-45% over 5 minutes, 45-97% over 2 minutes, hold for 1 min, and 97% to 3% over 2 min.

Full scan mass spectra (350–1800 m/z, 60,000 resolution) were detected in the orbitrap analyzer after accumulation of 3x10^6^ ions (normalized AGC target of 300%) or 25ms. For every full scan, MS/MS scans were collected during a 3 s cycle time. Ions were isolated (0.4 m/z isolation width) using the standard AGC target and automatic determination of maximum injection time, fragmented by HCD with 30% CE, and scanned at a resolution of 120,000. Charge states <2 and>4 were excluded, and precursors were excluded from selection for 30 s if fragmented n=2 times within a 20-s window.

### MS data search and manual inspection

All mass spectra were analyzed with Proteome Discoverer (PD, version 3.0) and searched using Sequest with rescoring using INFERYS and Percolator against a custom database comprising the Uniprot human proteome (UP000005640) together with the Uniprot *Mycobacterium tuberculosis* H37Rv proteome (UP000001584). No enzyme was used, and variable modifications included oxidized methionine for all analyses. Peptide-spectrum matches from MHC-II analyses were filtered with the following criteria: search engine rank = 1, length between 8 and 30 amino acids, XCorr ≥ 2.0, spectral angle ≥ 0.6, and percolator q-value < 0.05.

Identifications (IDs) of putative *Mtb*-derived peptides were rejected if any peptide-spectrum matches (PSMs) for the same peptide were found in the unfiltered DDA MS data for the corresponding mock-infected control. For each putative Mtb peptide identified, MS/MS spectra and extracted ion chromatograms (XIC) were manually inspected, and the ID was only accepted for further validation if it met the following criteria: (1) MS/MS spectra contained enough information to unambiguously assign a majority of the peptide sequence; (2) neutral losses were consistent with the chemical properties of the peptide; (3) manual de novo sequencing did not reveal an alternate peptide sequence that would explain a greater number of MS/MS spectrum peaks; (4) XIC showed a peak in MS intensity at the mass to charge ratio (m/z) of the peptide precursor ion at the retention time at which it was identified that did not appear in the corresponding mock-infected control. Peptides that met these criteria were further validated using SureQuant (see below).

### Synthetic standard survey MS analyses

Synthetic SIL standard peptides were synthesized by BioSynth as a crude peptide library. DDA MS analysis of the SIL peptide mixture was performed as described above (see DDA MS analyses) with the following modifications: Peptides were eluted using a flow rate of 300 nL/min and a gradient of 6–35% buffer B over 30 min, 35–45% over 2 min, 45–100% over 3 min, and 100% to 2% over 1 min. No dynamic exclusion was used.

A second set of survey analyses was performed on the mixture of SIL peptides with background matrix using the full SureQuant acquisition method (see below). SIL peptides were spiked into a sample of MHC-II peptides purified as described above from Raji cells, providing a representative background matrix. Because SIL amino acids are not 100% pure, SIL peptide concentrations were adjusted and survey analyses were repeated until the SIL peptide could be reliably detected while minimizing background signal detected at the mass of the biological peptide.

### SureQuant MS analyses

#### Validation analyses

Standard MS parameters and MS1 scan parameters were as described above (see DDA MS analyses), except that as none of the targets of these analyses has an m/z < 380, an MS1 scan range of m/z 380–1500 was used to exclude some common background ions. A flow rate of 300 nL/min was used. The custom SureQuant acquisition template available in Thermo Orbitrap Exploris Series 2.0 was used to build a validation analysis method targeting the set of putative *Mtb* peptides detected in DDA analyses for each donor. For each method, after the optimal charge state and most intense product ions were determined via a survey analysis of the synthetic SIL peptide standards (see above), one method branch was created for each m/z offset between the SIL peptide and biological peptide as previously described.^80^ SIL peptide precursor ions were matched with a mass accuracy tolerance of 10 ppm, and a threshold of n=3 out the top 6 product ions was used for pseudo-spectral matching with a mass accuracy tolerance of 20 ppm.

### Quantitative analyses

Quantitative SureQuant analyses were performed similarly, targeting hipMHC peptides in addition to biological peptides of interest. Data were analyzed using Skyline Daily Build 22.1.9.208. For each target peptide, the intensities of the three most intense product ions were integrated over the time during which the peptide was scanned. These intensities were normalized by the integrated intensities of the corresponding product ions from the corresponding SIL standard over the same time interval, and these ratios were averaged. Finally, these averaged light/heavy ratios were normalized by the average of the corresponding ratios for the hipMHC standard peptides and normalized to a reference condition (indicated for each experiment). 100 fmol of each SIL peptide was used, except AAGTYVAADAAAAST (EsxG; 1 pmol) and ASAAVEALTARLAAAH (PE5/PE29; 1 pmol).

#### Immunofluorescence microscopy

100,000 hMDCs per well plated on chamber slides (Ibidi) were infected with GFP-expressing *Mtb* at MOI 2.5 or mock-infected with media containing no Mtb. 72 hours post-infection, hMDCs were washed with PBS and fixed with 4% paraformaldehyde (PFA) in PBS for 1 hr. Slides were blocked with 5% normal goat serum in PBS supplemented with 0.3% v/v Triton X-100 for 1 hr at room temperature and stained overnight at 4 °C with primary antibody (LAMP1 – Cell Signaling Technologies [CST] D2D11; CTSB – BioLegend 948001) diluted in antibody dilution buffer (PBS with 1% w/v bovine serum albumin and 0.3% v/v triton X-100). Wells were washed three times with PBS for 5 min each and stained with Alexa fluor 647 (AF647)-conjugated goat anti-rabbit or anti-mouse secondary antibody (Thermo) diluted to a final concentration of 1 μg/mL in antibody dilution buffer at room temperature for 2 hr. Wells were washed three times with PBS for 5 min each and stained with DAPI at a final concentration of 300 nM in PBS for 15 min at room temperature. Wells were washed three times with PBS for 5 min each, well dividers were removed, and slide covers were mounted on slides using Prolong Diamond anti-fade mounting media (Thermo). Slides were imaged using a TissueFAXS Confocal slide scanner system (TissueGnostics). 25 fields of view were acquired per condition using a 40× objective lens.

#### Segmentation of Mtb-containing phagosomes

Image segmentation was performed using opencv-python 4.5.4.60. A two-dimensional Gaussian blur with a standard deviation of five pixels was applied to de-noise GFP fluorescence images. A binary mask was generated from the blurred GFP image using a fluorescence intensity threshold that was empirically selected for each biological replicate. GFP+ phagosomes were further segmented using the watershed algorithm.

#### Colocalization analysis

A correlation image was generated by computing the Pearson correlation coefficient between the GFP fluorescent intensity and AF647 fluorescent intensity in a 41x41 pixel sliding window with a step size of 1. This correlation value was averaged over each Mtb-containing phagosome, and phagosomes with a mean value of greater than or equal to 0.6 were considered co-localized.

#### MHC-II flow cytometry

A total of 1x10^6^ hMDMs or hMDCs were differentiated as described above on six-well plates. Where indicated, cells were infected with GFP-expressing *Mtb* at an MOI of 2.5 (or mock-infected with media containing no *Mtb*). 72 hours post-infection, phagocytes were detached by incubating in PBS supplemented with 4 mM EDTA for 15 minutes and gently dislodged with a cell scraper. Cells were blocked with Human TruStain FcX Fc receptor blocking solution (BioLegend) for 10 min, stained with phycoerythrin (PE)-conjugated anti-HLA-A,B,C antibody (BioLegend, clone W6/32) or PE-conjugated anti-HLA-DQ antibody (BioLegend, clone HLA-DQ1) and PerCP-Cy5.5-conjugated (non-*Mtb* experiments) or APC-conjugated (*Mtb* experiments) anti-HLA-DR antibody (BioLegend, clone L243) for 20 min, stained with Live/Dead Fixable Aqua dye (Thermo) or Zombie Violet dye (BioLegend) for 10 min, washed with FACS buffer and PBS, fixed with PBS containing 4% PFA for 1 hr, and washed and resuspended in FACS buffer for analysis on an LSR Fortessa flow cytometer (BD).

#### Analysis of mutations in MHC-II antigens

The collection of whole-genome sequences from 51,229 Mtb isolates were described previously.^39^ We used a previously described analytical pipeline to call SNPs of each isolate.^39^ Briefly, sequencing reads were trimmed with *Sickle*.^81^ Trimmed reads with length > 30 and Phred scores > 20 were retained for subsequent analyses. The inferred ancestral genome of the most recent common ancestor of the *MTB*C was used as the reference template for reads mapping.^82^ Sequencing reads were mapped to the reference genome using *Bowtie* 2 (v2.2.9)^83^ *SAMtools* (version 1.3.1) was used for SNP calling with the minimal mapping quality set to be 30.^84^ We excluded SNPs located in regions of the genome that are difficult to characterize with short-read sequencing technologies.^85^ Fixed SNPs with a frequency of ≥95% and at least 10 supporting reads were identified using *VarScan* (v2.3.9) with the strand bias filter on. *Mtb* isolates were typed into lineages based on the previously defined lineage-specific barcode SNPs.^86,87^

To define the dN/dS ratio for IEDB reported *Mtb* T cell epitopes, previously reported T cell epitopes were downloaded from IEDB (https://www.iedb.org/) in March 2023, using previously described criteria.^88^ Specifically, we applied the following filters: linear peptides, *M. tuberculosis* complex, positive assays only, T cell assays, any MHC restriction, host: human, any disease, and any reference type. This resulted in the identification of 2141 epitopes. Epitopes found in source proteins of MHC-II peptides identified by MS in our study were then excluded. Alignments of sequences encoding MHC-II peptides, non-epitope-containing regions from MHC-II peptide source proteins, and T cell epitopes reported in IEDB, were used to calculate pairwise dN/dS ratios.^89^ Pairwise dN and dS values were calculated using the R package seqinr with the kaks function, drawing 1,000 samples per iteration from each comparison group for 100 iterations. To avoid having undetermined pairwise dN/dS values due to dN or dS being zero, a mean dN/dS value was calculated for each sequenced isolate by dividing its mean pairwise dN by its mean pairwise dS with respect to all other sequenced isolates. The statistical differences between the MHC-II peptides and non-epitope regions of antigens, or between MHC-II peptides and IEDB reported T cell epitopes, were assessed using Wilcoxon rank-sum tests with continuity correction implemented in R version 3.2.2.

#### Publicly available immunogenicity data

The proportion of the cytokine-producing T cell response in healthy IGRA+ individuals attributed to each antigen by Panda et al. was retrieved from published source data where available. Panda et al. report percentages for individual proteins only for a subset of the proteome, whereas a moving sum over a 5-gene window was reported for the whole genome. The moving sum was deconvolved using non-negative least squares regression as implemented in Scipy 1.14.0 to retrieve data for individual antigens where not otherwise reported.

#### Synthetic peptides for ELISpot assays

Peptides were synthesized at the MIT Biopolymers and Proteomics Lab using standard Fmoc chemistry using an Intavis model MultiPep peptide synthesizer with HATU activation and 5 μmol chemistry cycles. Starting resin used was Fmoc-Amide Resin (Applied Biosystems). Cleavage from resin and simultaneous amino acid side chain deprotection was accomplished using: trifluoroacetic acid (81.5% v/v); phenol (5% v/v); water (5% v/v); thioanisole (5% v/v); 1,2-ethanedithiol (2.5% v/v); 1% triisopropylsilane for 4 hr. Standard Fmoc amino acids were procured from NovaBiochem. Peptides were quality controlled by MSy and reverse phase chromatography using a Bruker MicroFlex MALDI-TOF and Agilent model 1100 HPLC system with a Vydac C18 column (300 Å, 5 μm, 2.1×150 mm2) at 300 μL/min monitoring at 210 and 280 nm with a trifluoroacetic acid/H_2_O/MeCN mobile phase survey gradient.

#### ELISpot assays

ELISpot assays were performed using PBMC samples from a subset of a previously described participant cohort.^90^ The samples were originally collected as follows: Household contacts (HHCs) of newly diagnosed active pulmonary TB cases were identified through public health surveillance at community health facilities in Addis Ababa, Ethiopia, and their demographic and medical history data were collected. Active pulmonary TB (index) cases with drug-susceptible TB were symptomatic individuals with acid-fast bacilli (AFB) sputum smear positive or a positive GeneXpert MTB/RIF (Cepheid, Sunnyvale, California) result and a positive culture for Mtb growth. HHCs were persons who shared the same home residence as the index case for ≥5 nights during the 30 days prior to the date of TB diagnosis of the index case and were enrolled no more than 3 months (mean: 18 days; range: 1-77 days) after the index case began TB treatment. All participants provided written informed consent to join the study, and enrolled individuals met the following inclusion criteria: ≥ 13 years of age at the time of enrollment, positive QuantiFERON TB Gold in Tube (QFT) result, seronegative for HIV antibodies, no previous history of diagnosis or treatment for active TB disease or LTBI, normal chest X-ray, and not pregnant. The BCG vaccination status of participants was determined by checking for the presence of a BCG scar. The study was approved by the Institutional Review Board at Emory University, USA, the AHRI/ALERT Ethics Review Committee, and the Ethiopian National Research Ethics Review Committee. For each assay described here, PBMCs from donors expressing the HLA allele associated with a given antigen were used (Supplementary Table 3).

PBMC samples were selected from 7 asymptomatic QFT+ individuals in the HHC cohort who expressed class II HLA alleles associated with each MHC-II peptide of interest (Supplementary table 3). 5 of the 7 samples were from donors who were previously BCG vaccinated, while two (donors 4 and 5, Figure 2f) were not. Cryopreserved PBMC were thawed in a 37°C water bath and transferred into a 15mL conical tube with pre-warmed10mL R10 (RPMI 1640 supplemented with 1% PenStrep, 1% HEPES, 1% L-Glutamine [100x], and 10% FBS). The cells were centrifuged at 900g for 5 minutes at room temperature, supernatant discarded, cell pellet dislodged, transferred to a 6-well plate with 5ml R10, and rested for 3-4 hours at 37°C/5% CO2. Cells were counted based on Trypan blue exclusion criteria. Cells were resuspended in CTL-Test Medium (ImmunoSPOT-supplemented with 1% fresh L-Glutamine) and transferred to a 96-well plate at 200,000 cells/well in 100L volume. Mtb antigen peptides were added at a concentration of 2g/mL. Stimulated cells were cultured for 2 days at 37°C/5% CO2. For control, triplicate wells containing the same number of cells without antigen stimulation were included for each participant.

ELISPOT plates were activated according to manufacturer instructions (ImmunoSPOT) and human IFNγ /IL-17 capture solution was added and rested overnight at 4°C. The ELISPOT plate was then washed according to the manufacturer’s instructions, pre-stimulated cells were transferred from the the 96-well culture plate to the ELISPOT plate. Cells were cultured for an additional 24 hours (for a total of 72-hour stimulation). The ELISPOT plate was developed according to the manufacturer’s protocol. The developed plate was left to dry and read on the ELISPOT machine (S6 Ultra M2 – ImmunoSPOT).

<u>Data Cleaning:</u> Data was cleaned and manually checked and corrected as needed. Response magnitudes were determined after subtraction of background, which was calculated as the mean spot forming unit (SFU) of no-stimulation wells plus 2 times the standard error of the mean of the no-stimulation wells.

### Other statistical analyses

Unless otherwise indicated, all statistical tests of MS and ELISpot data were performed in GraphPad Prism 10.

## Data availability

Raw mass spectrometry data have been deposited to the ProteomeXchange Consortium via the PRIDE with the dataset identifier PXD055952.

## Acknowledgements

The authors thank all the staff members of the Ragon Institute, the Koch Institute, MIT, UCSF, NCI, and UNC for the essential work they do to make our research possible. We thank Lauren Stopfer, Cameron Flower, Tigist Tamir, Elizabeth Choe, Ryuhjin Ahn, and Melissa Bernhardt for helpful conversations, training, and technical guidance. We modified code written by Cameron Flower to plot MS/MS spectra. The Tü39 hybridoma used for in-house antibody production was generously provided by Hans-Georg Rammensee and Claudia Falkenburger. Alex Austin and Heather Amoroso (Koch Institute Biopolymers Core) synthesized, purified, and characterized peptides for ELISpot assays and MHC class II hipMHC production. MHC class II hipMHC complexes were produced by ImmunAware (Hørsholm, Denmark). This work is supported by funding from the MIT Center for Precision Cancer Research at the Koch Institute and U.S. NIH grants U01 CA238720, U54 CA283114, 1R35GM142900, R01A1022553, R01 AI173002, and U19 AI111211. O.L. is supported by U.S. NIEHS grant T32-ES007020. This work was performed in part in the Ragon Institute BSL3 core facility, which is supported by the NIH-funded Harvard University Center for AIDS Research (P30 AI060354). We thank Yong Xie, Julie Boucau, and Eliane Shwairi for managing the facility. This project has been funded in whole or in part with federal funds from the Frederick National Laboratory for Cancer Research, under Contract No. 75N91019D00024. The content of this publication does not necessarily reflect the views or policies of the Department of Health and Human Services, nor does mention of trade names, commercial products, or organizations imply endorsement by the U.S. Government. This Research was supported in part by the Intramural Research Program of the NIH, Frederick National Lab, Center for Cancer Research.

---

**Supplementary figure 1.**
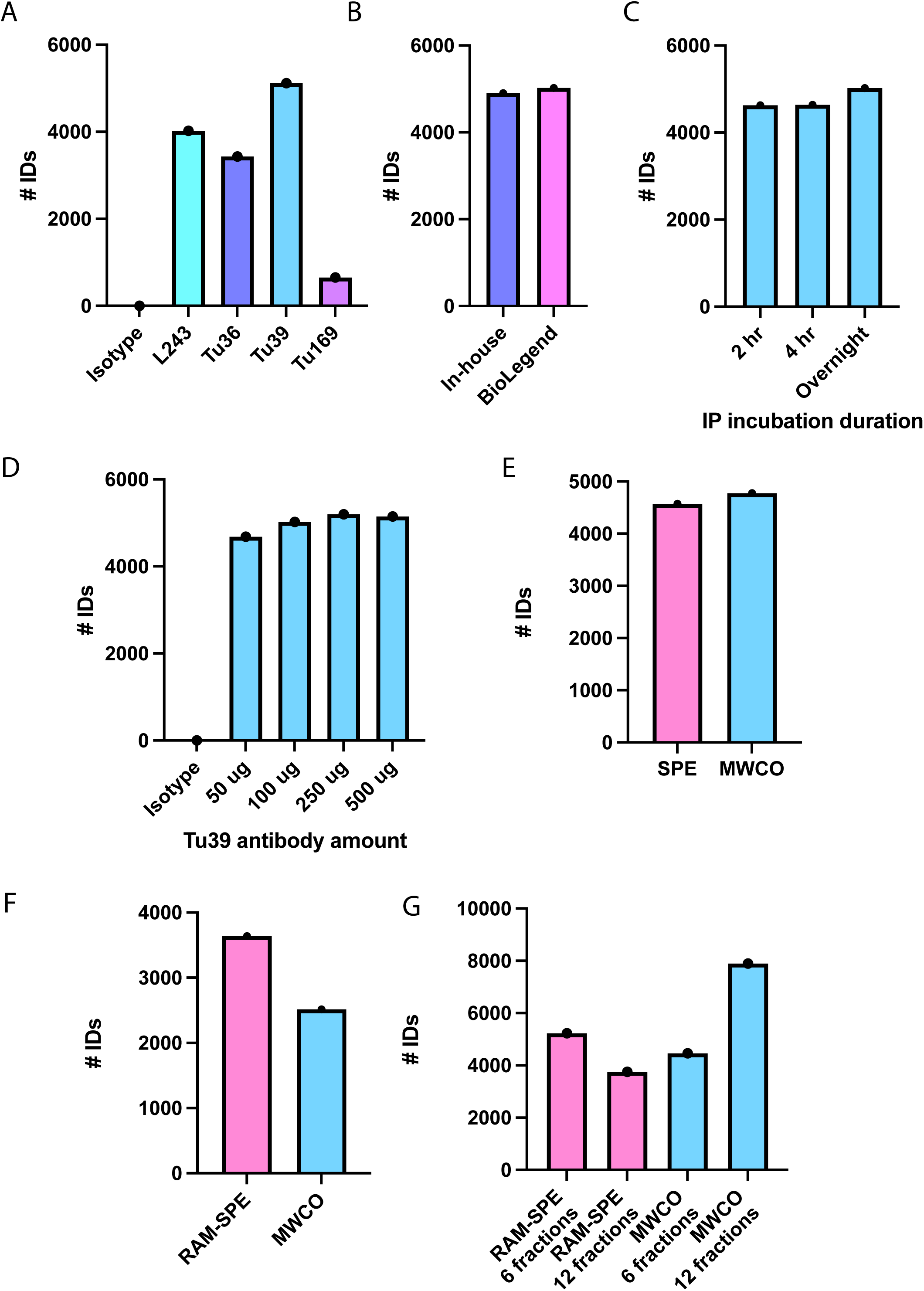
Optimization of a human MHC-II immunopeptidomics workflow. Number of unique peptide IDs obtained in analyses of the MHC-II immunopeptidome of Raji cells. Unless otherwise noted, IPs were performed using 100 μg of Tü39 antibody overnight (12-14 hours), peptides were isolated using a 10 kDa molecular weight cutoff (MWCO) filter, and analyzed by LC-MS/MS without prior offline fractionation. a) IPs performed using 100 μg of each of four MHC-II-specific antibodies. b) IPs performed using 100 μg of Tü39 antibody produced in-house or commercially produced. c) IPs incubated for 2 hours, 4 hours, or overnight. d) IPs performed using varying amounts of Tü39 antibody. e) MHC-II peptides isolated via solid-phase extraction (SPE) using a C18 spintip (Protea) or a MWCO filter. f) Peptides isolated using an in-house packed restricted access material solid phase extraction (RAM-SPE) column^79^ or a MWCO filter, without offline fractionation. g) MHC-II peptides prepared using either a RAM-SPE column or a MWCO filter and separated by offline reversed-phase HPLC into 60 1.5-minute fractions concatenated into 12 or 6 tubes (see Methods) prior to LC-MS/MS analysis.

**Supplementary figure 2.**
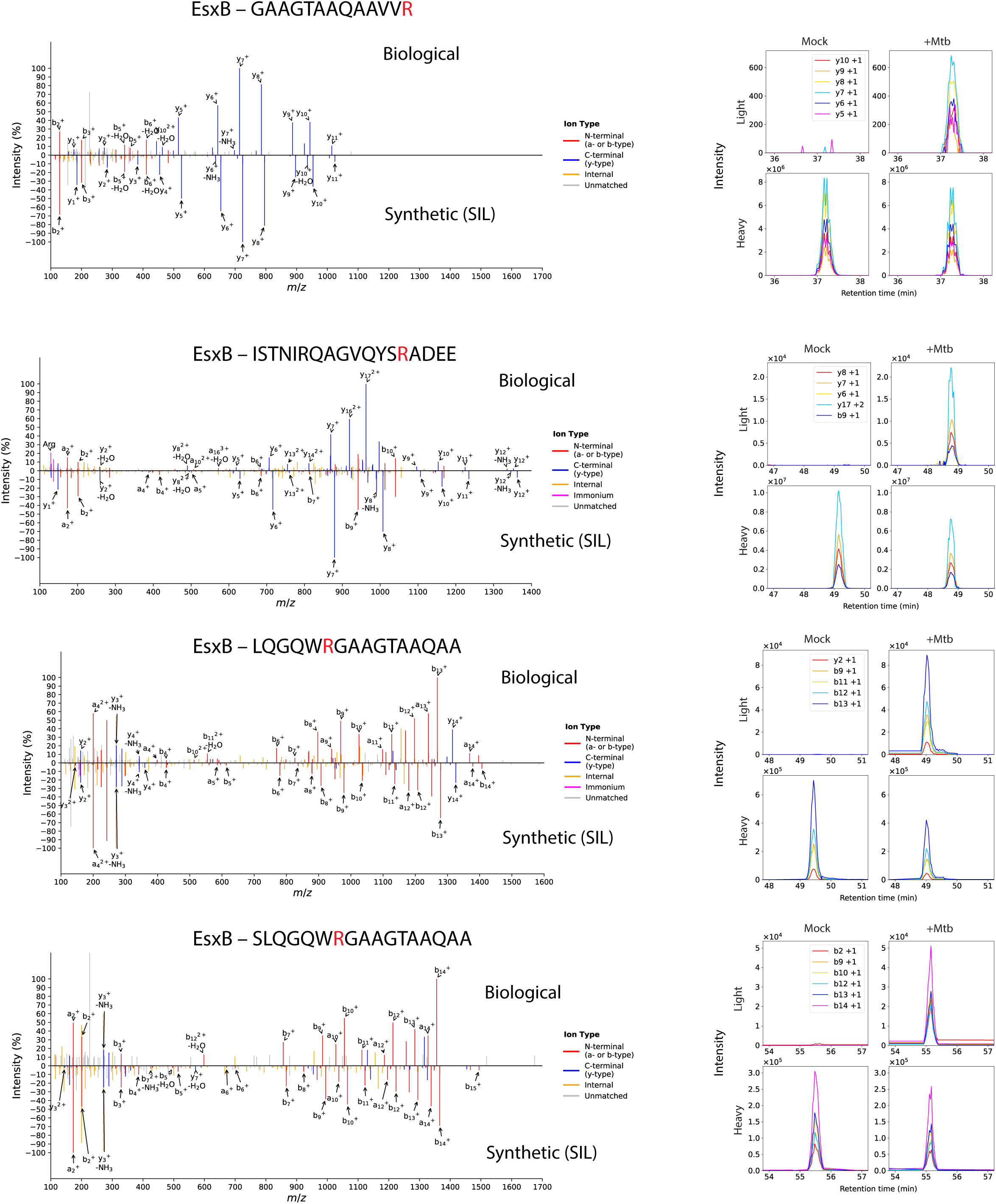
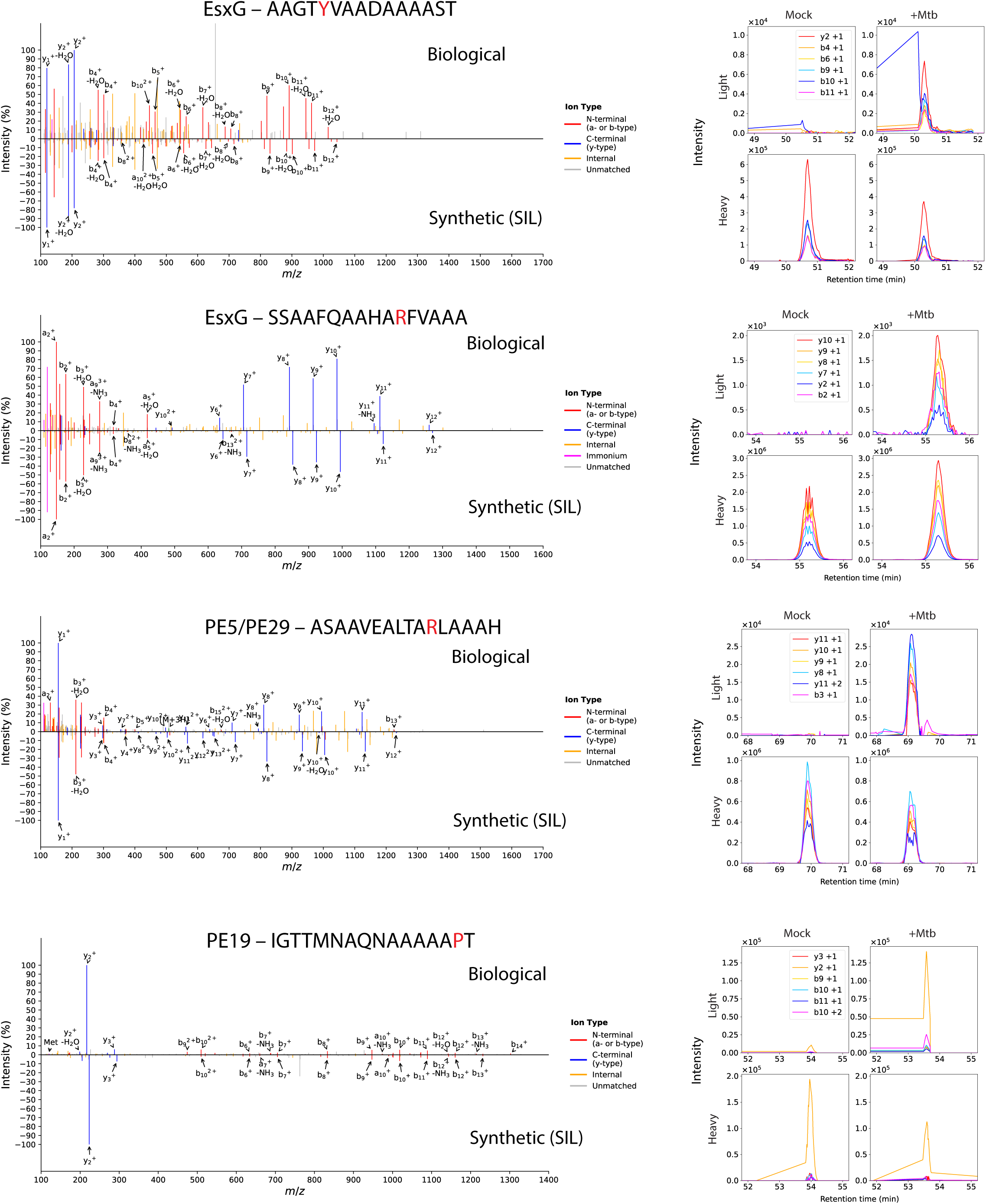
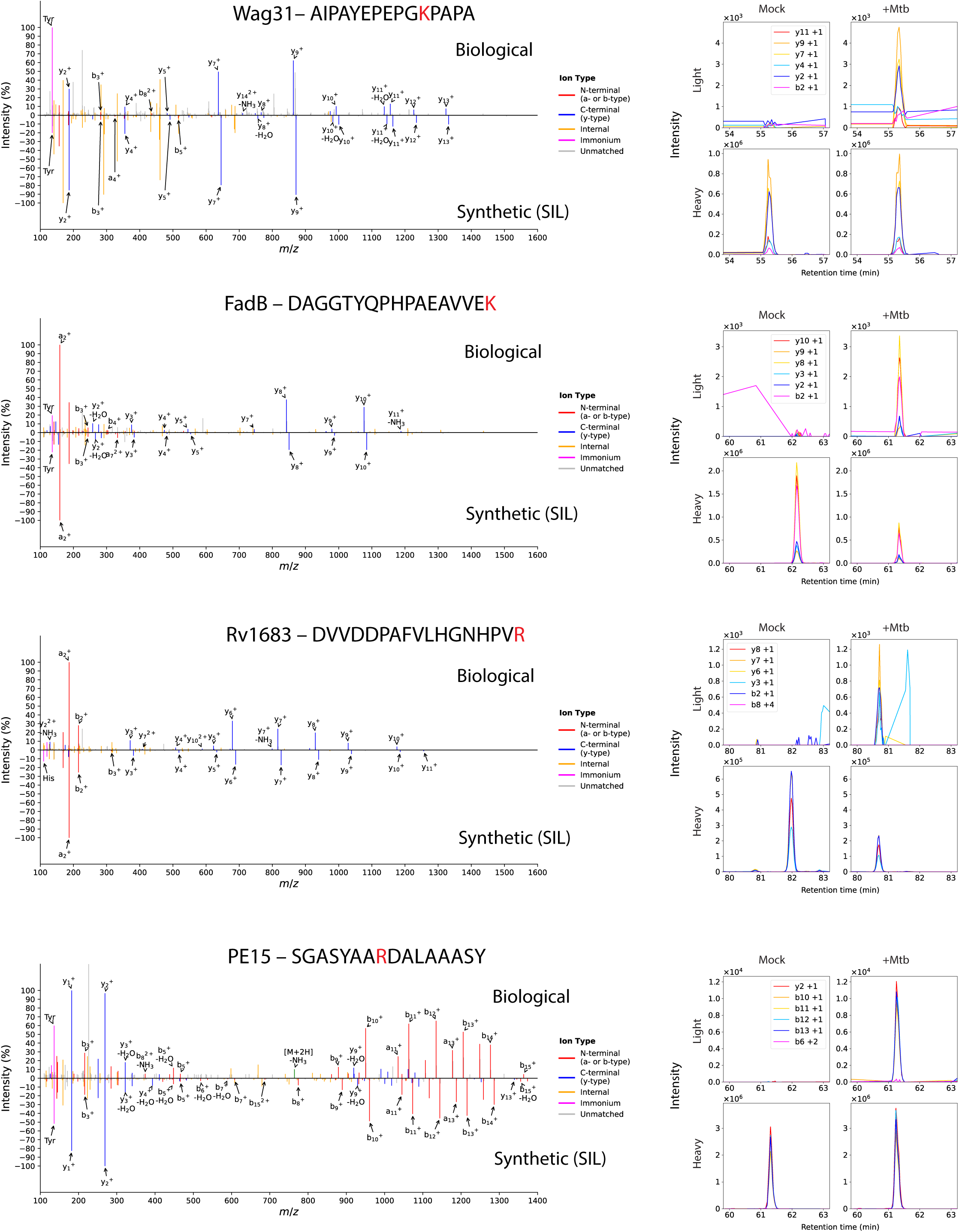
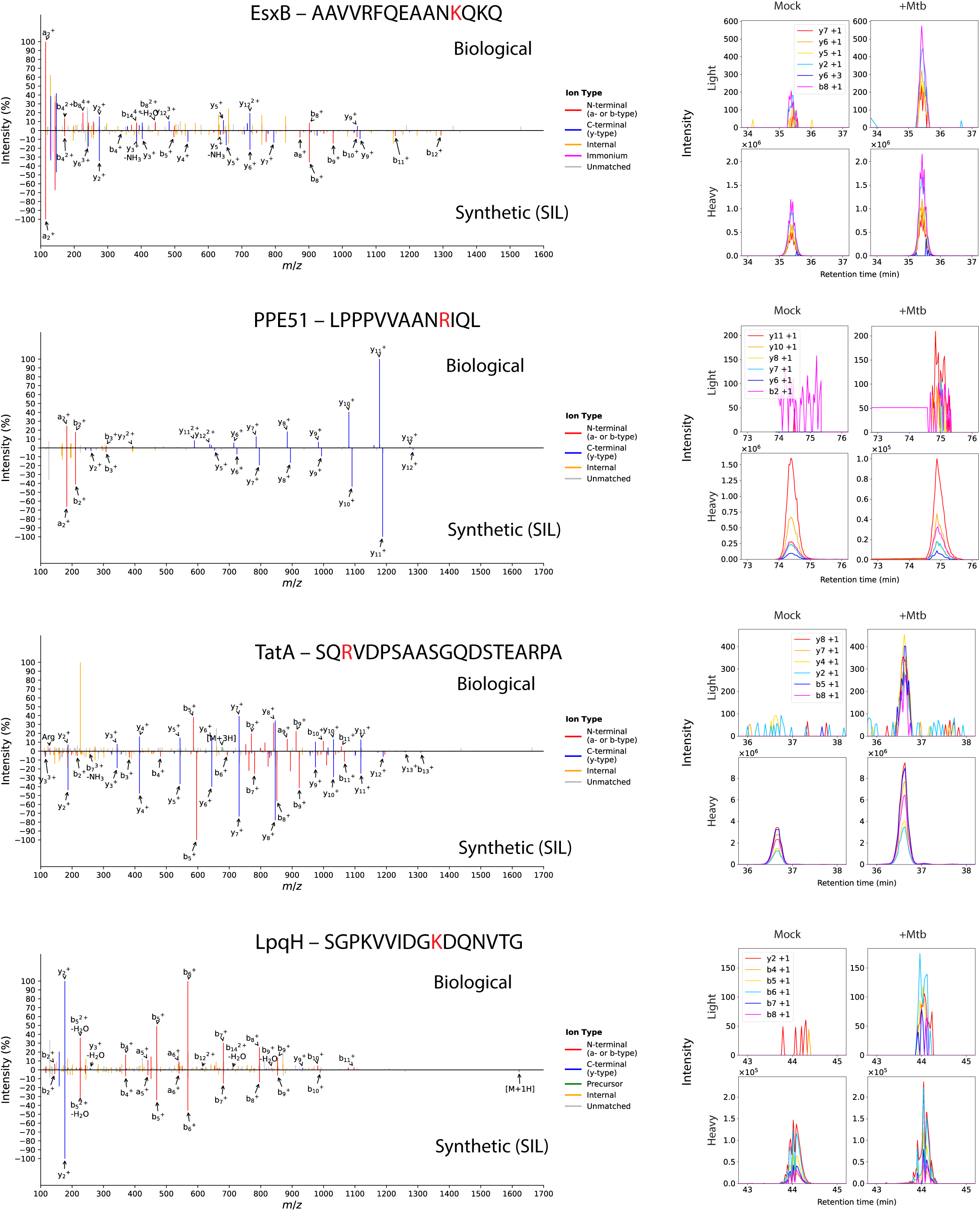
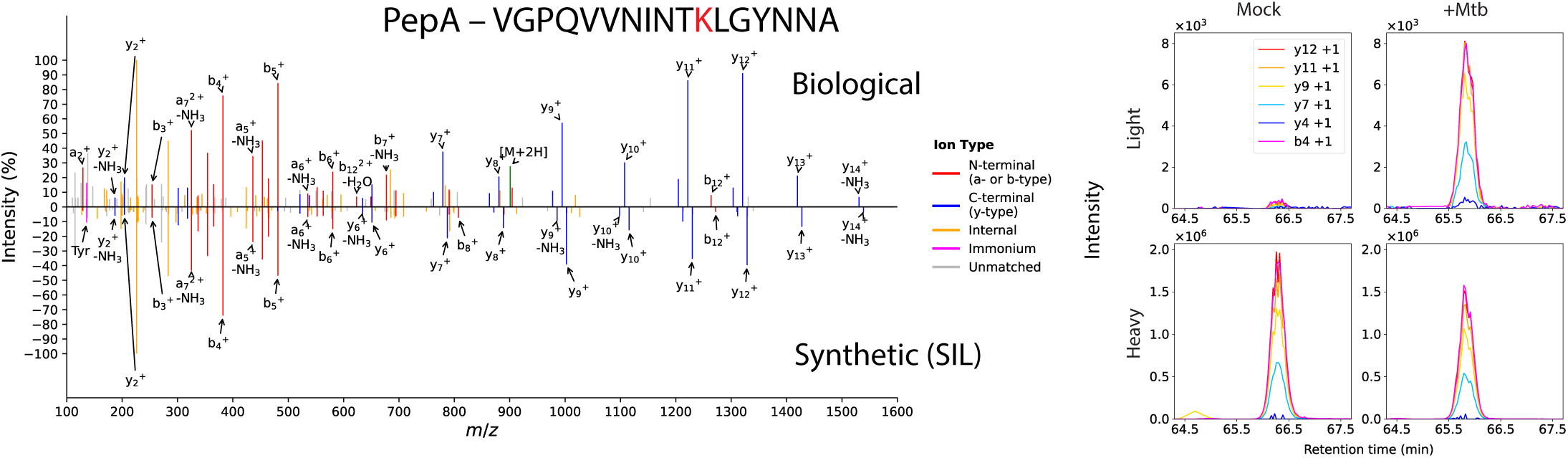
Validation of *Mtb*-derived MHC-II peptides by SureQuant. MS/MS spectrum comparisons (left) and SureQuant product ion chromatograms for *Mtb*-infected and mock-infected hMDCs (right column) for each peptide. For each peptide, the amino acid that is stable isotope labeled in the synthetic standard is indicated in red.

**Supplementary figure 3.**
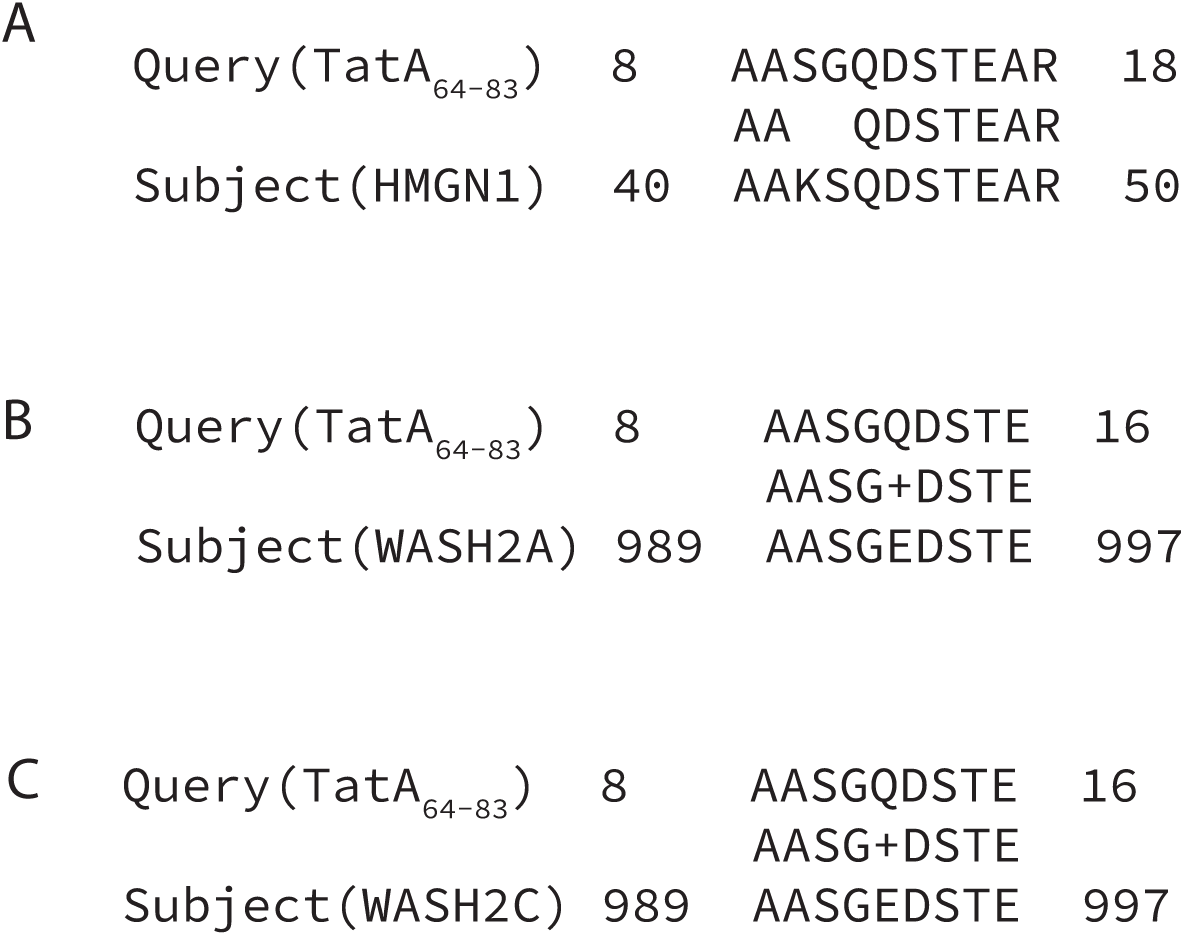
A TatA-derived MHC-II peptide shares sequence identity with human proteins. Uniprot BLAST search results using TatA_64-83_ as a query against the human proteome, showing regions of sequence identity between TatA_64-83_ and (a) HMGN1, (b) WASH2A, and (c) WASH2C.

**Supplementary figure 4.**
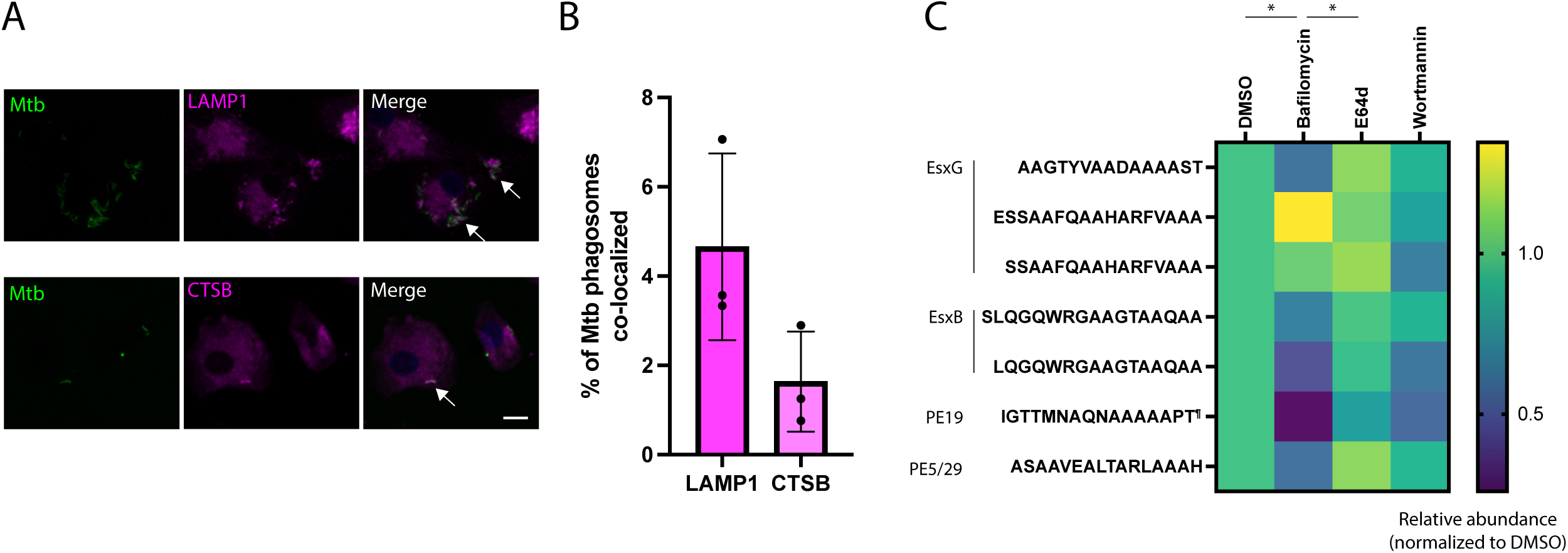
Interaction of *Mtb* with acidified endomembrane compartments may influence MHC-II antigen processing. a) Representative examples of co-localization of *Mtb* with LAMP1 and Cathepsin B (CTSB) in infected hMDCs visualized by immunofluorescence microscopy (IF). b) Quantification of the percentage of *Mtb*-containing phagosomes co-localized with LAMP1 and Cathepsin B (see Methods). c) Quantification of *Mtb*-derived peptides presented on MHC-II by SureQuant in HLA-DRB1*01:01-expressing hMDCs treated with bafilomycin, E64d, wortmannin, or DMSO alone. Heatmap values represent the mean of n = 3 donors. ^¶^This peptide was quantified in either the oxidized or non-oxidized state, depending on successful detection for each donor. (* p < 0.05, two-way main effects ANOVA with Tukey’s multiple comparisons test).

**Supplementary figure 5.**
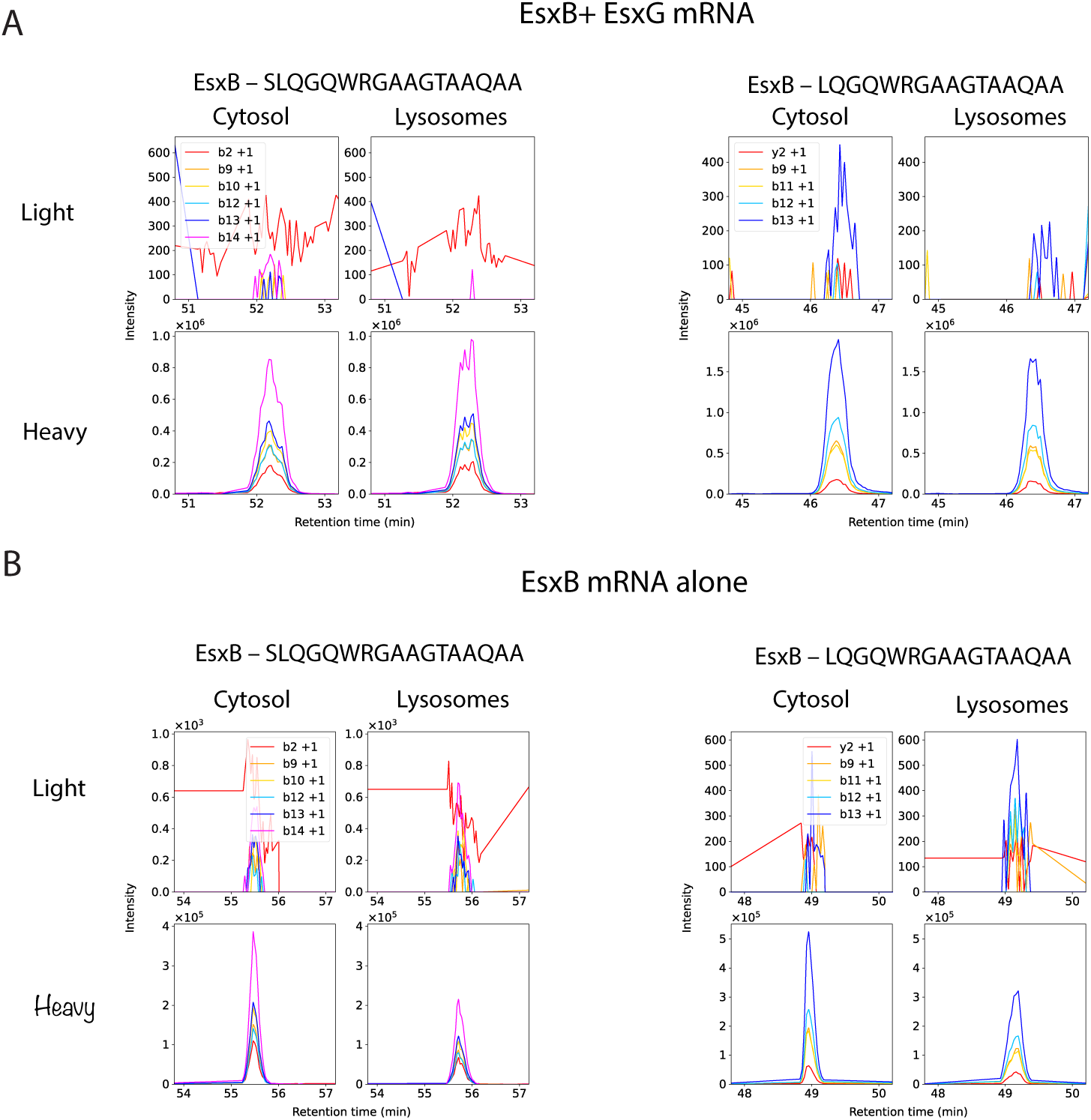
Presentation of EsxB-derived MHC-II epitopes is inefficient in hMDCs transfected with mRNAs encoding EsxB alone or EsxB and EsxG. Chromatograms of fragment ions for EsxB-derived peptides in MHC-II peptide samples from hMDCs transfected with mRNAs encoding EsxB and EsxG (a; representative of n = 3 donors) or EsxB alone (b; n = 1) fused to targeting signals for the cytosol or lysosomes.

**Supplementary table 1.** Parameters of prototype and final MHC-II immunopeptidomics workflows used in *Mtb*-infected cells.

| Donor(s) | Cell type | Number of cells | Antibody | Offline fractionation? (Y/N) | Gradient | Total unique peptide IDs (+*Mtb* condition) |
| --- | --- | --- | --- | --- | --- | --- |
| A | hMDC | 50 million | L243 | Y | Original MHC gradient<sup>18</sup> | 2619 |
| B | hMDC | 100 million | Tü39 | N | Original MHC gradient | 3345 |
| C | hMDM | 75 million | Tü39 | Y | Optimized MHC-II gradient (see Methods) | 1392 |
| D,E,F (final workflow) | hMDC | 50 million | Tü39 | Y | Optimized MHC-II gradient | 2700, 2877, 4390 |

**Supplementary table 2.** HLA-II genotypes of primary monocyte donors for immunopeptidomics.

| Donor | DPA1 allele 1 | DPA1 allele 2 | DPB1 allele 1 | DPB1 allele 2 | DQA1 allele 1 | DQA1 allele 2 | DQB1 allele 1 | DQB1 allele 2 | DRB1 allele 1 | DRB1 allele 2 | DRB3 allele 1 | DRB3 allele 2 | DRB4 allele 1 | DRB4 allele 2 | DRB5 allele 1 | DRB5 allele 2 |
| --- | --- | --- | --- | --- | --- | --- | --- | --- | --- | --- | --- | --- | --- | --- | --- | --- |
| A | 01:03 | 02:01 | 04:01 | 10:01 | 01:03 | 01:02 | 05:03 | 05:02 | 13:01 | 16:01 | 01:01 |  |  |  |  | 02:02 |
| B | 01:03 | 01:03 | 02:01 | 04:01 | 03:01 | 05:05 | 03:02 | 03:01 | 04:04 | 11:01 |  | 02:02 | 01:01 |  |  |  |
| C | 01:03 | 01:03 | 04:01 | 04:01 | 02:01 | 01:02 | 02:02 | 06:02 | 07:01 | 15:01 |  |  | 01:01 |  |  | 01:01 |
| D | 01:03 | 01:03 | 02:01 | 02:01 | 01:02 | 05:05 | 06:04 | 03:01 | 13:02 | 13:03 | 03:01 | 01:01 |  |  |  |  |
| E | 01:03 | 01:03 | 04:01 | 04:01 | 01:01 | 05:01 | 05:01 | 02:01 | 01:03 | 03:01 |  | 01:01 |  |  |  |  |
| F | 01:03 | 02:01 | 04:01 | 10:01 | 01:01 | 01:04 | 05:01 | 05:03 | 01:01 | 14:54 |  | 02:02 |  |  |  |  |

**Supplementary table 3.**
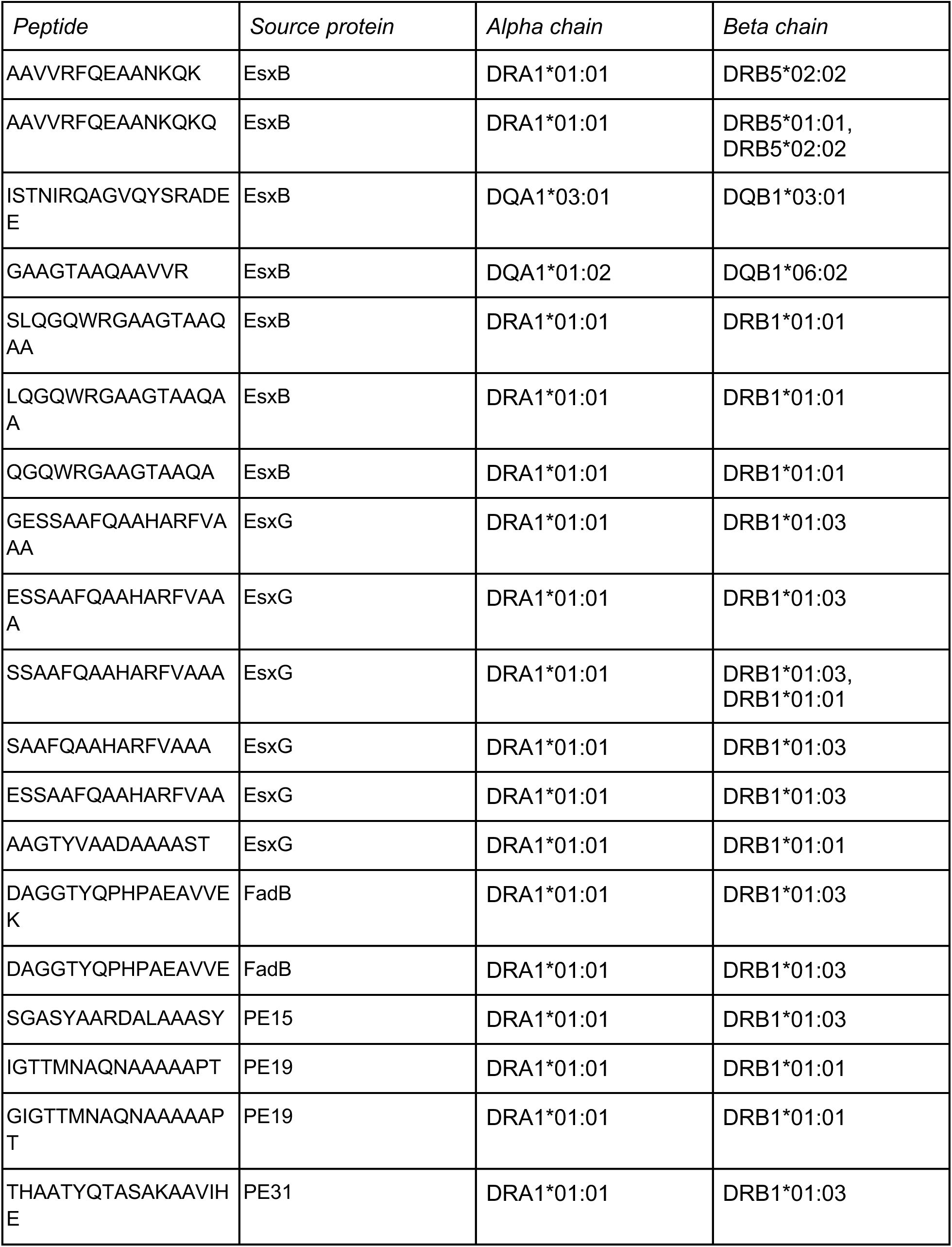
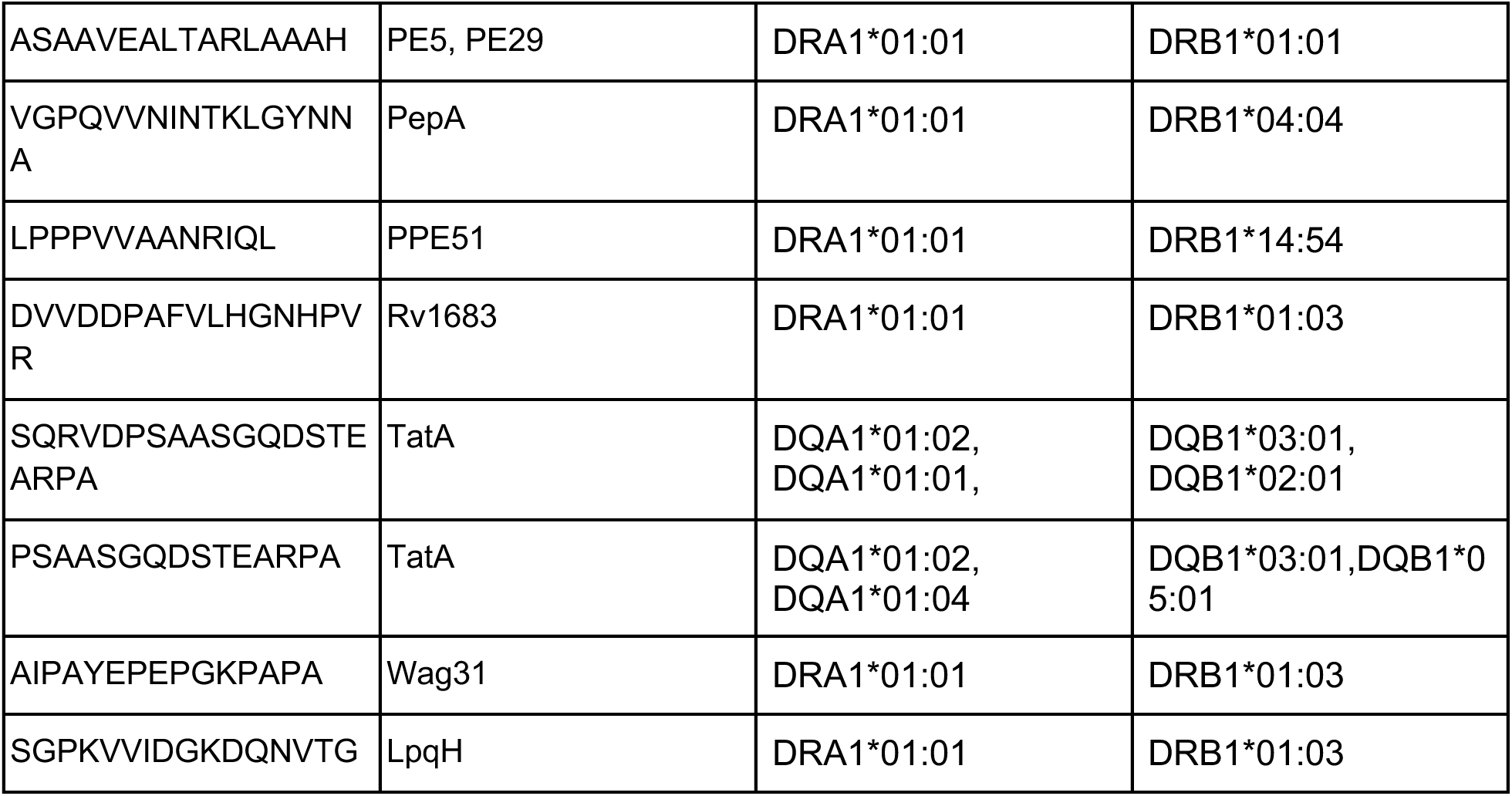
HLA-II alleles associated with *Mtb* MHC-II peptides (highest NetMHCIIpan predicted binding score, per donor)

**Supplementary table 4.** Sequence conservation of non-RD1 MHC-II epitopes and source proteins between BCG Pasteur and *Mtb* H37Rv.

| *Protein* | *MHC-II epitopes identity (%)* | *Whole protein identity (%)* |
| --- | --- | --- |
| EsxG | 100 | 100 |
| PE15 | 100 | 100 |
| PE31 | 100 | 100 |
| PE19 | 100 | 100 |
| PPE51 | 100 | 100 |
| PE5 | 100 | 100 |
| PE29 | 100 | 100 |
| LpqH | 100 | 100 |
| FadB | 100 | 99.86111111111111 |
| Rv1683 | 100 | 100 |
| TatA | 100 | 100 |
| PepA | 100 | 100 |

**Supplementary table 5.**
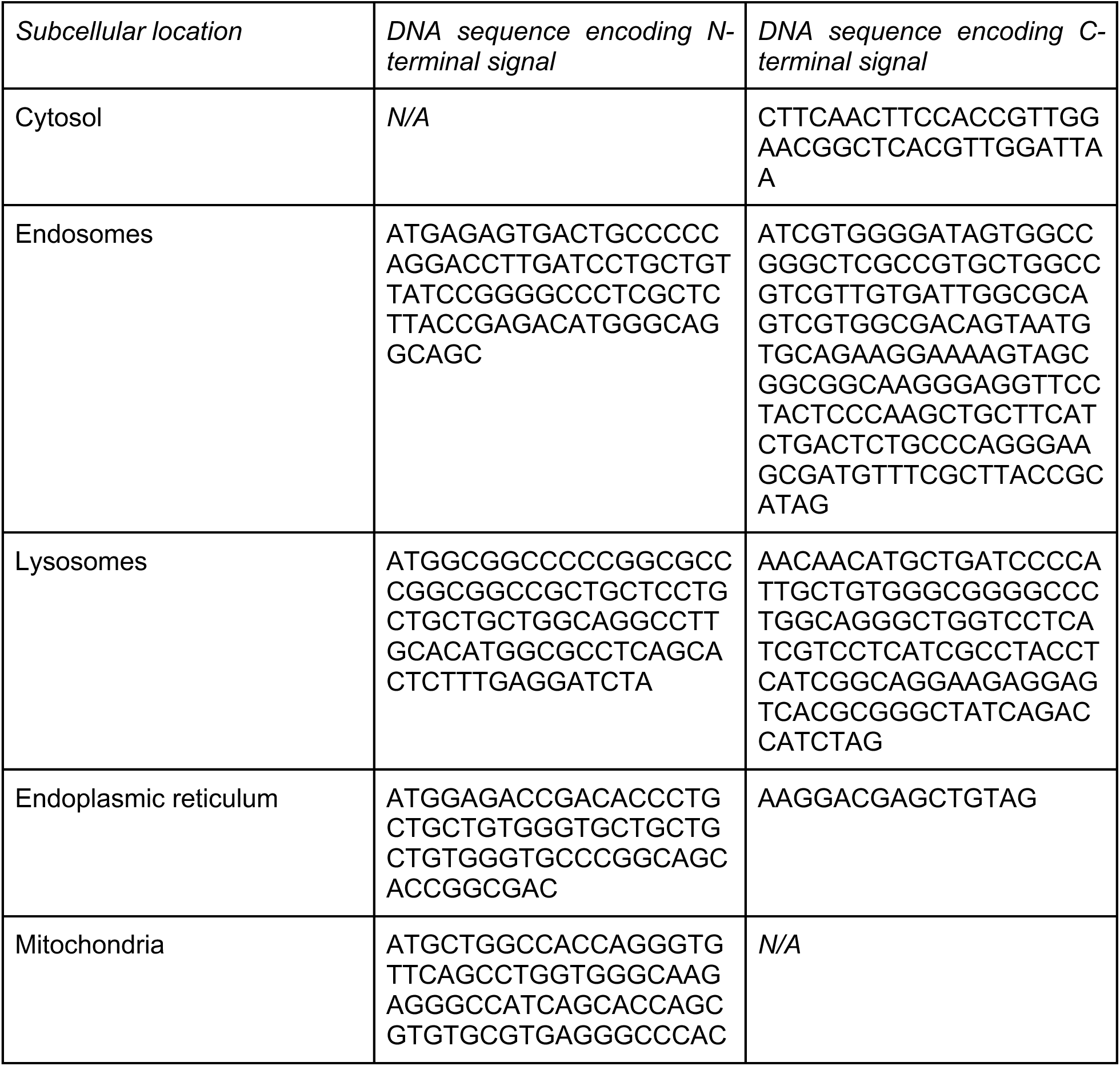
Subcellular localization signal sequences used in mRNA constructs.

Supplementary data 1. Analysis of mutations in *Mtb* genomic loci encoding MHC-II peptides

Supplementary data 2. Nonsense mutations upstream of *Mtb* MHC-II peptides

